# Characterising the Photoplethysmography Pulse Waveform for Use in Human Neuroscience: The Hybrid Excess and Decay (HED) Model

**DOI:** 10.1101/2021.08.19.456935

**Authors:** Simon Williamson, Lucie Daniel-Watanabe, Johanna Finnemann, Craig Powell, Adam Teed, Micah Allen, Martin Paulus, Sahib S. Khalsa, Paul C. Fletcher

## Abstract

Photoplethysmography (PPG) offers a widely-used, convenient and non-invasive approach to monitoring basic indices of cardiovascular function such as heart rate and blood oxygenation. However, while the pulse waveform, generated by PPG comprises features that are shaped by physiological and psychological factors, it is frequently overlooked in analyses of such data. We suggest that studies could be enriched by exploiting the possibilities afforded by a systematic analysis of PPG waveforms. To do this we initially require a robust and automated means of characterising it, thereby allowing us to examine variations across individuals and between different physiological and psychological contexts. We present a psychophysiologically-relevant model, the Hybrid Excess and Decay (HED) Model, which characterises pulse wave morphology in terms of three underlying pressure waves and a decay function. We show that these parameters capture PPG data with a high degree of precision and, moreover, are sensitive to specific, physiologically-relevant changes within individuals. We present the theoretical and practical basis for the model and demonstrate its performance when applied to a pharmacological dataset of 105 participants receiving intravenous administrations of the sympathomimetic drug isoproterenol (Isoprenaline). We conclude by discussing the possible value in using the HED model to complement standard measures of PPG outputs.

## 1 INTRODUCTION

Photoplethysmography (PPG) is an inexpensive and non-invasive means of measuring heart rate and peripheral oxygen saturation. It is ubiquitous in healthcare settings, increasingly prevalent in wearable devices (Zhang Y 2020) and of growing interest in cognitive neuroscience research as an indirect measure of autonomic arousal, its ease of use making it applicable at a large scale (Natarajan A 2020). It is also frequently employed to provide the objective data against which individuals’ subjective awareness of their own heartbeats can be compared i.e. as a measure of interoception (Khalsa, et al. 2008; Smith, et al. 2021). Here, we provide a technical examination of the PPG signal, and suggest that, complementing the more frequently-derived measures such as heart rate (HR) and heart rate variability (HRV), its shape provides a rich, valuable, and often overlooked source of information. The haemodynamic events underlying this shape are complex and offer the potential for a more granular assessment of cardiovascular features which, in turn, are of relevance to physiological and psychological states. Thus, focusing on morphological features of the PPG signal could lead to new characterizations of psychophysiological states relevant to mental health and cognition.

The PPG signal is acquired using a paired light emitter and sensor. The sensor measures the quantity of emitted light that returns after transmission or reflection through the tissue. Changes in the composition of the tissue – such as fluctuations in local peripheral blood volume - are therefore reflected in the signal. Typically, the signal is described in terms of its alternating current (AC) and direct current (DC) components (Allen, 2007; Akl *et al*., 2014). The AC component comprises the visible fluctuation in signal generated by arterial pulses, with each pulse giving rise to a distinct waveform spanning the length of the cardiac cycle. The DC component represents the light absorbed by venous blood and the remaining tissue. Fluctuations in the DC component are reflected in the signal baseline and are influenced by other physiological phenomena such as respiration, sympathetic nervous system activity, and thermoregulation (Allen, 2007).

The resulting waveform measured using PPG has a distinctive morphology (see figure 1) related to that of the central aortic pressure wave (Imholz *et al*., 1998). From the field of arterial haemodynamics, there are two distinct but complementary models for understanding how the waveform is shaped:

**Figure 1.**
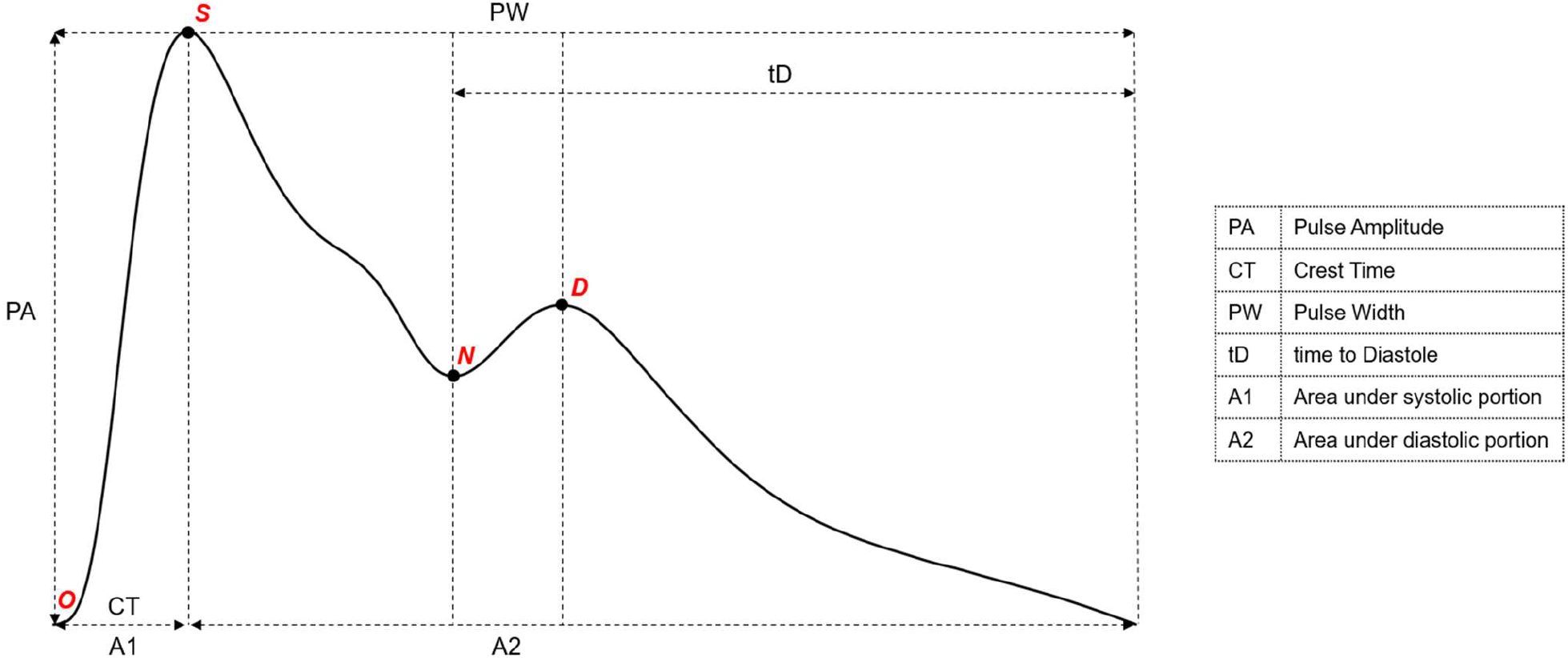
The Pulse Waveform and features identified as being of psychophysiological relevance. The predominant fiducial points, O, S, N and D, represent the Origin, Systolic peak, Dicrotic Notch and Diastolic peak of the waveform, respectively (Elgendi, Liang and Ward, 2018). Derivative measures found to be of psychophysiological relevance are labeled; PA = pulse amplitude (Khandoker *et al*., 2017); tD = time to Diastole (Rinkevičius *et al*., 2019); CT = crest time; PW = Pulse Width; A1 = area under the systolic portion of the waveform; A2 = area under the diastolic portion of the waveform (Charlton *et al*., 2018).

i. *The Windkessel Model:* Cardiac systole generates a pressure wave that is sustained by the elastic compliance of the arterial vessels, resulting in an initial surge followed by a slowly decaying pressure wave over the diastolic period (Westerhof, Lankhaar and Westerhof, 2009).
ii. *The Wave Model:* The initial systolic ejection is followed by one or more reflectance waves arising from distal sites in the arterial tree where impedance to flow is greater (van de Vosse and Stergiopulos, 2011). The precise nature and origin of these reflectance waves is not fully understood (Westerhof *et al*., 2008) but they have an appreciable effect on the overall shape of the wave and may vary within and between individuals in ways that are highly informative.

Both models suggest that the waveform is highly sensitive to cardiovascular state and therefore that its shape is rich in information about this state. Given the interest in psychological and physiological factors influencing, and potentially influenced by, cardiovascular state (Penninx, 2017; Mulcahy *et al*., 2019), a comprehensive characterisation of the features of this waveform could be a powerful complement to standard measures of heart rate and heart rate variability in characterizing and monitoring psychophysiological states. The work we describe here aimed at achieving this and, as we show, combines elements of both the above models.

Existing work has demonstrated the value of the pulse waveform as a marker for physiologically meaningful and clinically relevant information (Allen, 2007). The majority of features have been identified in the time domain through contour analysis: the identification of fiducial points from higher order derivatives (Millasseau *et al*., 2006; Elgendi, 2012). Additional approaches include frequency domain (Nitzan *et al*., 1994), non-linear (Sviridova and Sakai, 2015), and machine learning methods (Zhang *et al*., 2021). By these means a number of associations between cardiovascular factors and the waveform have been identified. These factors include blood pressure (Elgendi *et al*., 2019), central arterial stiffness (Millasseau *et al*., 2002), systemic vascular resistance (Lee *et al*., 2011), blood lipid (Sattar *et al*., 2014) and glucose levels (Asekar, 2018), and response to vasoactive drugs (Takazawa *et al*., 1998). Importantly for our purposes, changes in psychological state are also detectable in the waveform due to associated autonomic and cardiovascular effects (Abbod *et al*., 2011; Charlton *et al*., 2018; Rinkevičius *et al*., 2019) (see figure 1).

There are perhaps grounds for believing that the waveform offers a window onto processes highly relevant to brain-body interaction insofar as it contains information that the brain could use via vascular afferents to gain access to the peripheral body state. This would subsequently be useful in computing actions for goal-directed behavior or to augment exteroceptive input with interoceptive afferent information to inform a more adaptive response. An approach based on a more complete analysis of the waveform (rather than one that is focused on pre-selected points of interest) will allow for inferences to be made about the underlying haemodynamics and the factors affecting these.

Pulse decomposition analysis (PDA) operates on the understanding that the waveform is composed of several waves, and that these can be inferred. It offers the potential for waveform changes, which may be driven or shaped by relevant psychophysiological processes and hence are psychologically relevant, to be characterized in greater detail, that is, in terms of the component waveforms and their inter-relations. In modelling each waveform, the entire wave contour is considered and can be expressed in terms of the parameters used to generate the component waves. In addition, PDA-derived measures correlate closely with contour-based measures whilst demonstrating greater robustness to noise (Fleischhauer *et al*., 2020). In the development of these models, a question remains as to the optimal number of composite waves and which kernels are used to represent them.

Here, we present a prototype for characterising the PPG waveform: the Hybrid Excess and Decay (HED) model. It is comparable to PDA insofar as it expresses the waveform and its variability in terms of its component waves but it involves the addition of three further parameters to characterise the quasi-exponential decay following systole. We describe and present the model along with an automated means of applying it to PPG data. We go on to demonstrate its utility by applying it to data from a study in which the bolus effects of intravenous isoproterenol, a potent beta-adrenergic agonist, were compared to those of saline in a sample of 118 participants (from which 105 datasets were suitable for study). We assess the model’s performance, in terms of goodness-of-fit, on this PPG dataset as well as its ability to recapitulate the key fiducial points seen in figure 1. We then assess its ability to distinguish isoproterenol and saline conditions in order to examine its comparability to standard measures of physiological response (heart rate and heart rate variability) as well as existing established contour analysis measures. We conclude by considering the model’s concurrence with existing concepts in haemodynamics, and its potential for optimizing the usefulness of the PPG signal in characterizing psychological and physiological states.

## 2 METHODS

Our approach comprised two main parts:

2.1 *Model development:* This entailed (i) A theoretical analysis of the most likely constituents of the PPG waveform and (ii) refinement of the model and its assumptions with the use of a small pilot dataset.
2.2 *Model validation:* An analysis pipeline was developed to test the model on a larger dataset acquired on different hardware from that used in the pilot dataset. This dataset came from a study in which cardiovascular parameters were systematically manipulated using isoproterenol infusion. We aimed to examine (i) how well the model performed across both the drug and placebo infusions and (ii) its sensitivity to the pharmacological challenge (isoproterenol versus saline) in comparison to other standard measures, including heart rate, derived from PPG.

### 2.1 Model Development

#### 2.1.1 Theoretical Foundation and Model Components

A single prevailing view on the nature of arterial waves and their reflections has not yet been reached, largely due to the inherent complexity of the arterial tree (Segers *et al*., 2017). Nonetheless it is generally accepted that backward travelling reflectance waves exist and are likely to be measurable as composites from multiple reflection sites, irrespective of origin. Hitherto, three component waves of the pulse waveform have been described:

- A systolic wave caused by the increase in pressure, and therefore volume, arising from left ventricular contraction and ejection of oxygenated blood into the arterial system.
- A first reflectance wave, also described as the ‘renal’ wave (Baruch *et al*., 2011).
- A second reflectance wave, often referred to as the ‘diastolic’ wave (Millasseau *et al*., 2006).

PDA models have taken a largely data-driven approach, using combinations of component waves of varying number and shape. Comparisons of these suggest three to be the optimal number for capturing the waveform’s morphology (Tigges *et al*., 2017). Our model started from this point, modelling the waveform as a composite of three component waves (systolic, first reflectance wave (R1) and second reflectance wave (R2) as above). Nine parameters characterised the timing, width and amplitude of each component wave.

Our model, however, differed from existing PDA models in its approach to the diastolic decay. We noted a tendency for waves of prolonged duration to decay below baseline, an established phenomenon in aortic pressure waveform studies (Parker, 2017). To our knowledge this is not explicitly dealt with by current PDA models. To account for this and better elucidate the underlying factors driving waveform morphology, we incorporated a further three parameters:

- Decay rate: The rate at which the signal decays exponentially to baseline in the absence of component wave influence (Wang *et al*., 2003).
- Baseline 1: The baseline towards which the signal decays during the systolic portion of the waveform.
- Baseline 2: The baseline towards which the signal decays during the diastolic portion of the waveform.

In summary, therefore, we modelled the waveform as a composite of signal due to the initial systolic pressure wave, an exponential decay, and two reflectance waves. For simplicity, we refer to the systolic and reflectance waves as the “excess” elements and the exponential decay as the decay element (see figure 2), referring to the overall model as the Hybrid Excess and Decay (HED) model. The output of the model is a 12-parameter vector, which can be used to construct a fitted wave to the original PPG data (see figure 2).

**Figure 2.**
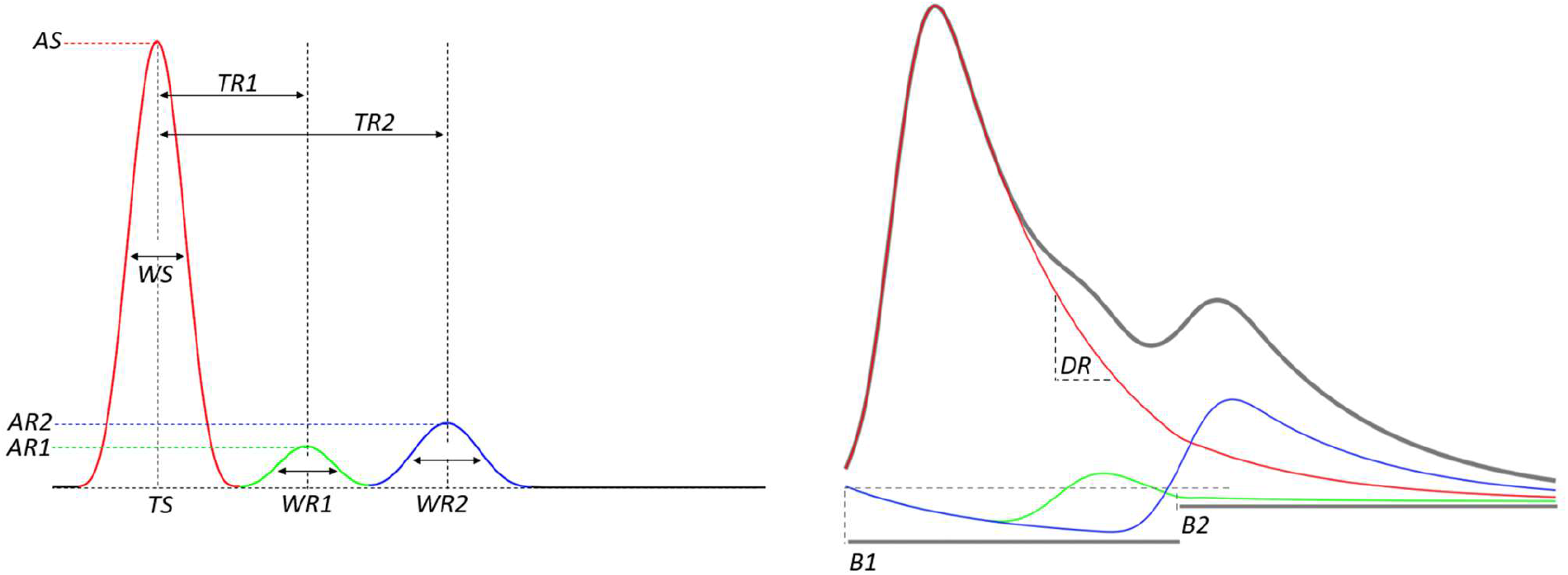
2a: The first 9 parameters of the model compose the excess element. *TS = Timing of systolic wave; TR1 = Timing of 1st reflectance wave; TR2 = Timing of 2nd reflectance wave; AS = Amplitude of systolic wave; AR1 = Amplitude of 1st reflectance wave; AR2 = Amplitude of 2nd reflectance wave; WS = Width of systolic wave; WR1 = Width of 1st reflectance wave; WR2 = Width of 2nd reflectance wave*. 2b: The final 3 parameters compose the decay element, which when incorporated with the excess yield the final modelled waveform. *DR = Decay rate; B1 = 1st baseline; B2 = 2nd baseline*.

#### 2.1.2 Refinement of the Model and Assumptions

Once established conceptually, the model was refined using a set of test data samples acquired from members of the experimental team at rest using a Bioradio device (Great Lakes Neurotechnologies, Ohio, USA); a multichannel portable device designed for simultaneous acquisition of multiple physiological signals (glneurotech, 2020). The accompanying PPG sensors are transmission-based finger clips made of either hard or soft plastic. Both types were used at the index finger, with a sampling rate of 75 Hertz (Hz). This small test dataset served as the basis for iterative assessment and correction of aberrant model behaviour, as described in the following paragraphs.

As is common in PDA (Fleischhauer *et al*., 2020), a number of assumptions were made in developing the model. For the excess components, sine waves were used as they exhibit a number of suitable properties: first, a sine wave smoothly interpolates to zero which fits with the appearance of PPG waves, insofar as the wave onset is not sudden; second, we found it useful to look at the first derivative of the PPG waveform in order to identify beats and this derivative is well-fitted by a sine wave. For the decay element, a second baseline spanning the diastolic portion was included to allow for more flexible modelling of varying dicrotic notch (see figure 1) height. Additionally, we assumed that (i) component waves cannot have the same timing and must occur in order, (ii) the 1^st^ reflectance wave (“renal wave”) plays a relatively minor role in shaping the waveform and must therefore have the smallest amplitude and width, and (iii) the number of component waves need not be three if a more parsimonious fit can be achieved with fewer. Constraints of the model reflect these assumptions and are detailed in full at: (https://github.com/Algebreaker/PulseWaveform).

Further, we noted that 6 of the 12 model parameters tended to converge on constant values across multiple resting heartbeats. These were (i) width of systolic wave (ii) width of 1st reflectance wave (ii) width of 2nd reflectance wave (iv) timing of 1st reflectance wave (v) timing of 2nd reflectance wave and (vi) decay rate. The option to fix these parameters across multiple beats during parameter optimization was added. Doing so greatly reduced the number of free parameters across time series (in brief, the number of free parameters was reduced by ((n/2 +6) where n is the number of free parameters (12) multiplied by the number of beats in the dataset)) as well as allowing greater consistency in how adjacent beats were fitted. There was a corresponding reduction in computational time required. We therefore maintain this function for the analyses herein but stress the optionality of this feature in our shared code such that waves can still be modelled individually if desired.

### 2.2 Model Validation

Formal testing of the model took place on a dataset from an ongoing functional magnetic resonance imaging (fMRI) project investigating the effect of adrenergic receptor stimulation on physiological, subjective, and neural measures of interoception (ClinicalTrials.gov Identifier: NCT02615119). This is hereafter referred to as the ISO study given its focus on the effects of isoproterenol (isoprenaline). For details regarding the development and implementation of this pharmacological perturbation approach, see (Khalsa et al 2009; Khalsa et al 2016; Hassanpour 2018). Participants in the study were predominantly female (93.3%; n=98) with a mean age of 24.2 (SD=5.7; range = 18-39). The sample was clinically heterogenous combining both healthy (n=45) and individuals with various psychiatric conditions (generalized anxiety disorder (n=29), major depressive disorder (n=7), and anorexia nervosa (n=24). The experimental procedure entailed the randomized administration of two bolus infusions of saline and two bolus doses of isoproterenol (2 micrograms (mcg) and 0.5 mcg, each dose repeated once) during fMRI scanning, though analyses undertaken here excluded data from the 0.5 mcg time series (rationale described in 2.2.2). Participants were blinded to the order of administration. Importantly, PPG data were collected using different hardware to that used to develop the model: GE MR750 MRI scanner equipment (i.e., hard plastic finger clip) affixed to the index finger of the nondominant hand and recorded at a sampling rate of 40 Hz while in the supine position.

### Summary of the Model Pipeline

The PPG processing pipeline employed for the ISO study analysis is laid out in figure 4 and comprised: (i) Pre-processing (ii) Beat segmentation (iii) Morphological feature extraction (contour analysis) (iv) Pulse decomposition modelling.

**Figure 3.**
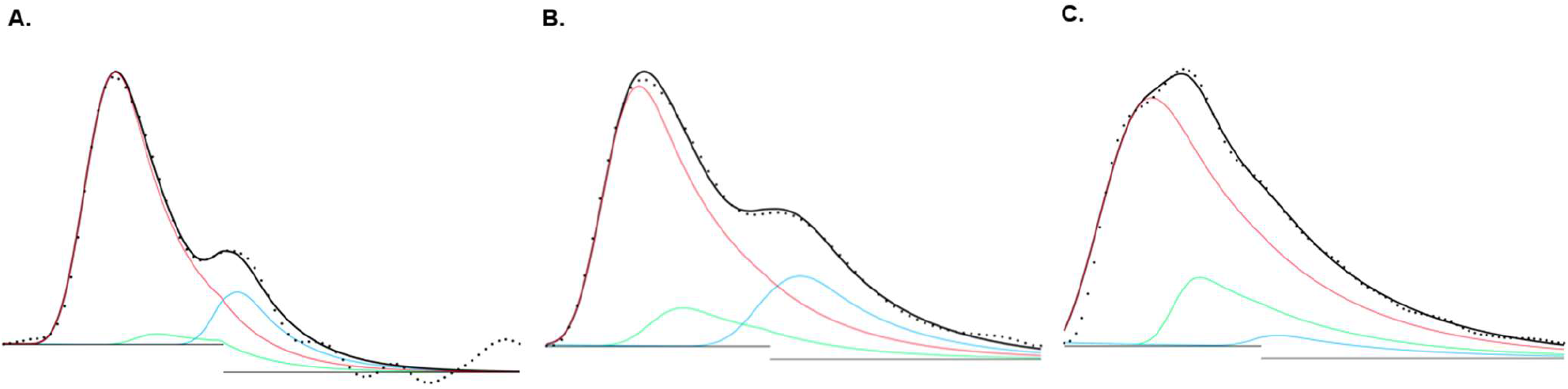
Three waves (acquired on Bioradio devise (see main text) fitted with the model. Each pulse wave was recorded from a different individual and is of a different morphological class (Dawber, Thomas and McNamara, 1973). a: class 1, Goodness of fit indicated by NRMSE = 0.93; b: class 2, NRMSE = 0.94; c: class 3, NRMSE = 0.85.

**Figure 4.**
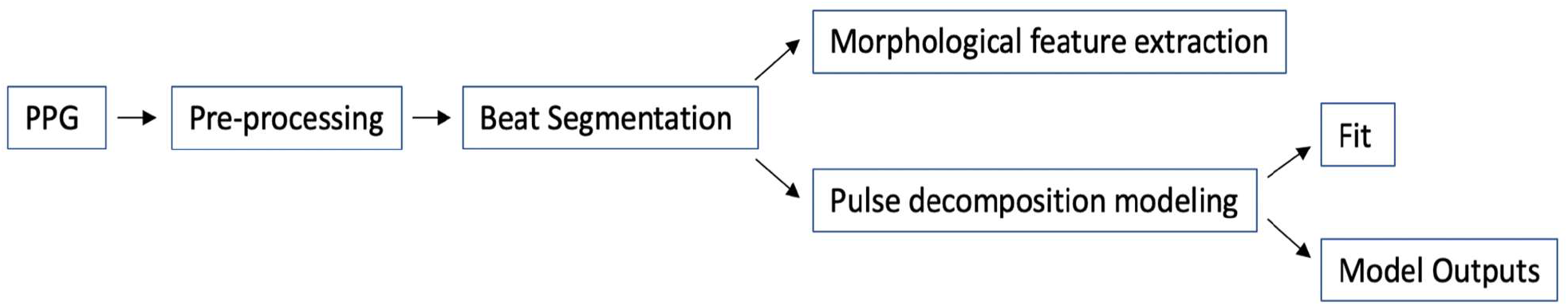
Figurative representation of the PPG data processing pipeline employed for the ISO test dataset. Open source code in R using *DescTools, pracma, splines2* and *SplineUtils*: (https://github.com/Algebreaker/PulseWaveform).

All beats underwent (iii) and (iv) with the former applied in order to generate a more “standard” analysis against which the performance of the model could be assessed. The output of the decomposition step was also assessed in terms of goodness of fit to the dataset as a whole (i.e. isoproterenol plus saline)

#### (i) Pre-processing

Many forms of PPG hardware automatically detrend raw data to maintain a constant baseline. This can confound contour analysis by altering the shape of the waveform. We employed an optional step to reverse the effect of detrending in two stages: (a) Adjusting the fixed amplitude drop with each successive datapoint such that the central portion of the waveform did not drop below baseline, and (b) Correcting for the change in gradient of the entire time series resulting from step (a).

#### (ii) Beat segmentation

A custom peak detection algorithm identified peaks in the first derivative of the time series (Elgendi, Liang and Ward, 2018) (sensitivity 0.97, positive predictive value 0.99, further details are provided in supplementary material (S1)). Identification of the first peak of this derivative (the so-called w peak) provided a marker of systolic waves and, thereby, a convenient means of calculating inter-beat intervals, and thereby, HR and HRV. The minima preceding *w*, denoted *O*, gave an indication of the wandering baseline of the signal, and through spline interpolation were used to remove the baseline drift (or DC component) from the time series. This ensured that only the arterial signal was assessed for any given waveform. *O* points also defined the start and end of each beat and were used to derive individual beat segments.

#### (iii) Morphological Feature Extraction

Each waveform was assessed in turn to derive fiducial points *O, S, N* and *D* (henceforth *OSND*) (see figure 1), with use of the first derivative. In the absence of a clear notch, the inflection point closest to zero in the first derivative was instead taken as N (Lee *et al*., 2011). To increase robustness against misidentification of points due to noise, *OSND* values for the average wave were identified first and informed identification on individual waves. Morphological features were then derived from *OSND* directly or with additional information from the segmented waveform (e.g., area under the wave). All waves were scaled for amplitude. Further derived measures are outlined in supplementary material (S2).

#### (iv) Pulse Decomposition Modelling

##### Initial parameter estimation

Initial parameter estimates were calculated per beat for the entire time series and were first estimated from the derived excess of a given waveform. In the excess, the systolic peak was tallest, followed by R2 and then R1. The 9 peak parameters could therefore be estimated sequentially. The 3 decay parameters were initially given fixed values.

##### Parameter Optimization

Initial parameter estimates were optimised for each waveform (and across waveforms - see 2.1.2). To achieve this we used the downhill simplex method (Nelder and Mead, 1965), which is a straightforward and robust means of finding the minimum of a function in multi-dimensional space (Press *et al*., 2007). In our case the function to optimize is goodness of fit, defined for each waveform as the reduced chi-squared (χ^2^*v*) is given by:

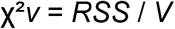

*Where RSS = The sum of squared (weighted) residuals, V = number of datapoints - number of parameters. Residuals are weighted such that a fitting of the central morphology of the waveform (from w to D) is prioritised*.

With no convention for the number of parameters used in PDA models, the model took an adaptive approach. χ ^2^*v*, by definition, penalizes non-parsimonious use of parameters. Since this function drives the optimization process, each waveform was fitted by the minimum number of parameters necessary to generate an optimal fit. In practice, this drove model behaviour such that, for a given waveform, two component waves were selected over three if goodness of fit was not significantly worsened by doing so. This approach is comparable to some existing PDA models (Wang et al, 2013).

The simplex was restarted to avoid convergence on local minima. Without an established means of assessing how many restarts are sufficient to converge on a global minimum, three restarts (four runs total) were chosen for simplicity and computational efficiency. Optimized parameters from the fourth run were taken as model outputs.

#### 2.2.1 Assessment of Model Performance

We aimed to demonstrate the external validity of our model by testing its performance on data from the ISO study (see above). Importantly, the hardware used to collect the PPG signal for that study was different from the hardware we used to develop the model in the pilot study, allowing an assessment of the generalisability of the model and associated processing pipeline. For each modelled waveform, goodness of fit was assessed. In the absence of a standard measure for goodness of fit in PDA, we use the normalized root mean square error (*NRMSE*) as reported by Sorelli *et al*., among others (see supplementary material – S3). A value of 1 indicates a perfect fit to the data, whilst a value of 0 indicates no improved performance over a null model (Sorelli, Perrella and Bocchi, 2018).

In a further assessment of the model’s performance, we used it to attempt to recapitulate standard morphological points modelled from each waveform in the form of *OSND* values. Model-generated waves were created and these values were extracted and compared to the corresponding values taken from the real data. This allowed an assessment of concordance between the model and measures of proven clinical relevance.

#### 2.2.2 Model Sensitivity to a Pharmacological Perturbation

During the ISO study, effects of bolus infusions of either saline or isoproterenol during PPG measurement were examined. Isoproterenol is a rapid-acting beta-adrenoceptor agonist similar to adrenaline that has been used for decades in psychophysiological research (Cleaveland CR 1972) (Contrada RJ 1991) (Cameron OG 2002) (S. Khalsa, et al. 2009). It targets β1 receptors, exerting a positive inotropic and chronotropic effect on the heart. It also equivalently targets β2 receptors, inducing peripheral vasodilation. By comparing the PPG signal during isoproterenol infusion with that acquired during saline, we aimed to test the waveform’s sensitivity to a cardiovascular perturbation and the model’s ability to characterize it.

After subjecting all data (across both isoproterenol and saline) to morphological and pulse decomposition analyses (see ‘summary of model pipeline’), we assessed the performance of features derived from both methods as classifiers between dose levels. We aimed to determine (i) if traditional morphological features are more sensitive to the pharmacological manipulation than heart rate or heart rate variability (ii) if model parameter outputs are more sensitive to the manipulation than either of the above. For simplicity of comparison and to ensure a reasonable cardiovascular effect size for characterization, PPG data corresponding to the 0.5 mcg dose level were not included.

The duration and subsequent effects of bolus isoproterenol infusions are short. Accordingly, we analysed the subset of the data comprising only beats from the infusion period of each time series and a matched sample from the saline infusion (for a summary of this process see supplementary material – S4). Given the risk of sampling bias in time series with few beats to analyse (Allen, 2007), sensitivity analyses were conducted excluding (i) no time series (ii) time series with fewer than 30 identified beats and (iii) time series with fewer than 60 identified beats.

## 3 RESULTS

We present our results in two parts:

3.1 *General Model Performance:* Performance on the ISO test dataset as assessed by (i) goodness of fit (*NRMSE)* (ii) Recapitulation of important fiducial points (*OSND*).
3.2 *Characterization of a Pharmacological Manipulation:* The assessment of model-derived and standard PPG metrics as classifiers between dose levels.

The ISO study dataset comprised 118 human participants. Of these, 6 were excluded due to obtrusive clipping of the PPG signal, 1 for missing data, and 3 for suspected mislabelling. An additional 3 were excluded on running the pipeline due to excessively noisy signals. Thus, a total of 105 participants were included in the following analysis. Each participant had two time series available at each dose level, both of which were included. The mean number of beats sampled was 67 (+/-25), ranging from 12 to 161. Results presented below were consistent across sensitivity analyses.

### 3.1 General Model Performance

#### 3.1.1 ISO data (General Model Fit)

Waves in the dataset were robustly fitted with an average (median) NRMSE value of 0.9 obtained across participants (figure 5), comparable to previously reported models (Wang *et al*., 2013; Sorelli, Perrella and Bocchi, 2018). See supplementary material figure S1 for alternative NRMSE values).

**Figure 5.**
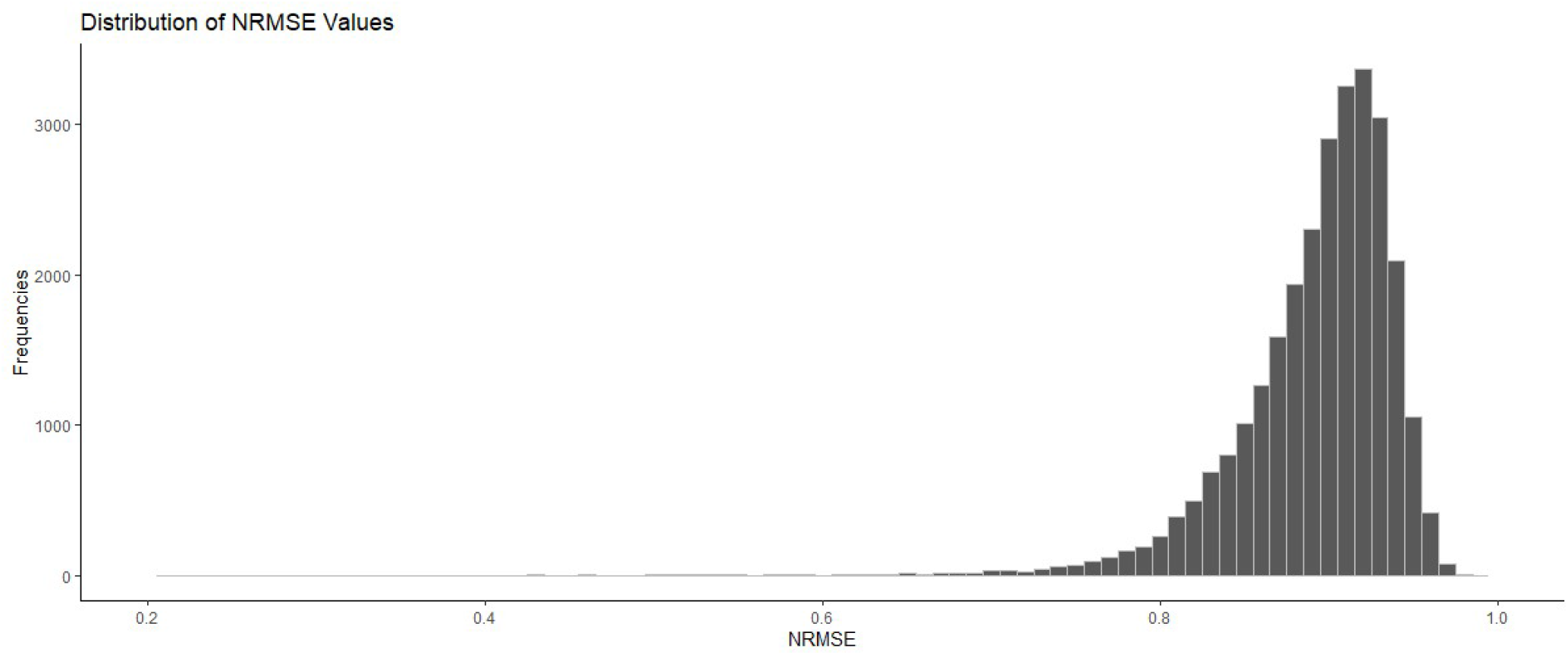
Distribution of NRMSE values for all waves fitted with the proposed model (binwidth=0.01). (For goodness of fit expressed in terms of residual error – *a*NRMSE see supplementary material (S3 and figure S1).

#### 3.1.2 ISO data (OSND Fit)

Figure 6 shows the accuracy of the model in regenerating fiducial points *OSND*. In order to illustrate this, a representative waveform and corresponding *OSND* values are plotted. Around each point model error (in the form of 65% and 95% confidence boundaries) is represented by ellipses. A tendency for the model to displace both *O* and *S* vertically is apparent, as well as *N* and *D* diagonally.

**Figure 6.**
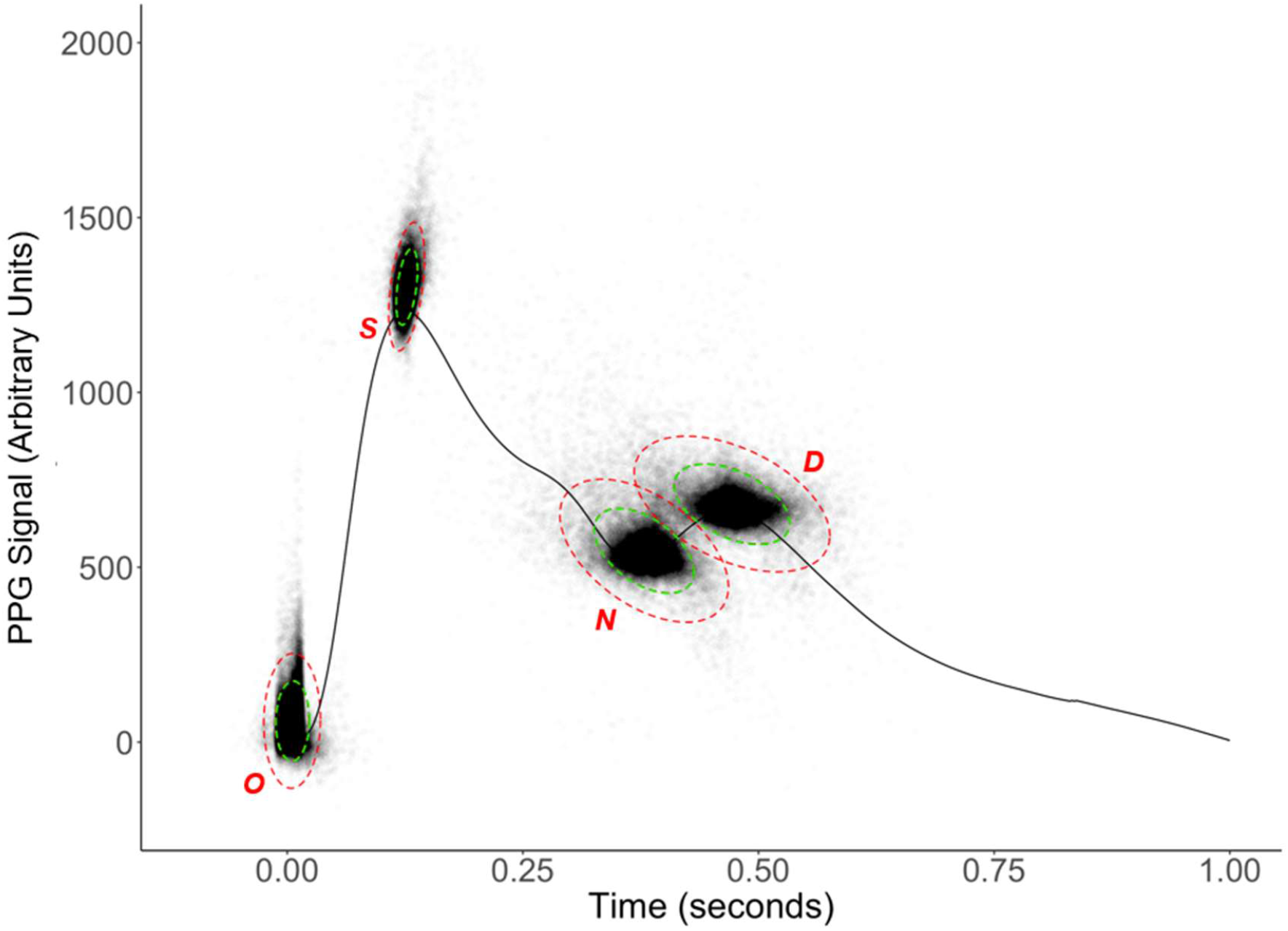
The distribution of error in the model’s fitting of key fiducial points *O, S, N* and *D*. Around each point model error (in the form of 65% and 95% confidence boundary) is represented by ellipses. A multivariate normal distribution is assumed.

### 3.2 Ability of HED model to characterise the effects of isoproteronol

For illustrative purposes, figure 7 shows the average waves for each of the 105 participants’ time series (from the subset of data related to the relevant periods of the isoproteronol or saline infusion) as well as an overall average across participants. These are presented separately for the isoproterenol and saline infusions. There appeared to be greater variability in morphology between participants at the saline dose. We therefore used the Levene test to compare homogeneity of variance between dose levels. This was carried out at each time point between w (corresponding to the first peak of the first derivative marking the systolic rise) and the end of the wave. This region of analysis lasted 0.865 seconds and consisted of 346 time points. For an alpha level of 0.05, the Bonferroni corrected p-value was 0.00014. The variance significantly differed across isoproterenol and saline a continuous period of 0.425 seconds (from 0.233 seconds to 0.658 seconds after w, which corresponds approximately to the peak of R1 and the end of R2.

**Figure 7.**
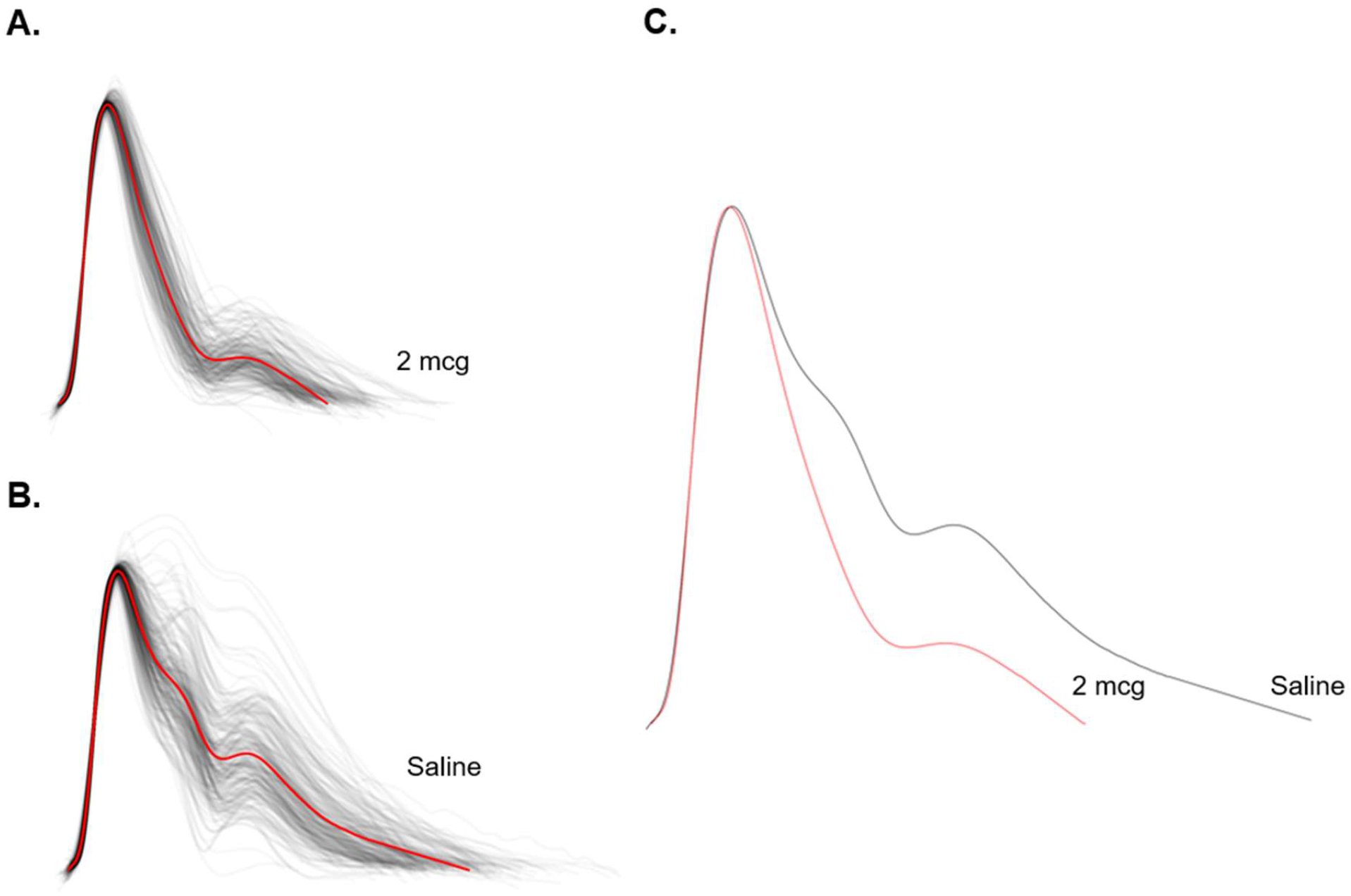
a: Averaged waveforms (black) during pharmacological manipulation (one per subject/time series) aligned by systolic peak. The mean of these (red) is overlaid. b: Averaged waveforms during infusion of saline (one per subject/time series) aligned by systolic peak. The mean is overlaid in red. c: The mean waves derived from a and b, overlaid to display the morphological change between saline and 2 mcg isoproterenol infusion.

Aside from this difference in variance within conditions, importantly, there were striking morphological differences between drug and saline (see figure 7 c). The isoproterenol (2 mcg) waveform shows a significantly lowered diastolic portion, with a steepened systolic decay and shortened diastolic decay. The overall length of the waveforms differs, as would be expected with an adrenergically-mediated increase in heart rate.

Differences in morphology between dose levels were first quantified using standard measures including fiducial point measures relating to contour analyses as well as heart rate and heart rate variability (HRV). SDNN was selected as the measure of HRV since a number of beats had to be rejected due to noise. Consequently, measures such as RMSSD, which are dependent on distances between successive beats, were considered sub-optimal. Those that characterized the difference most effectively are included in figure 8. Consistent with figure 7 c, these measures are directly related to either height of the diastolic portion (N amplitude, D amplitude), heart rate (wavelength) or both (area under the curve).

**Figure 8.**
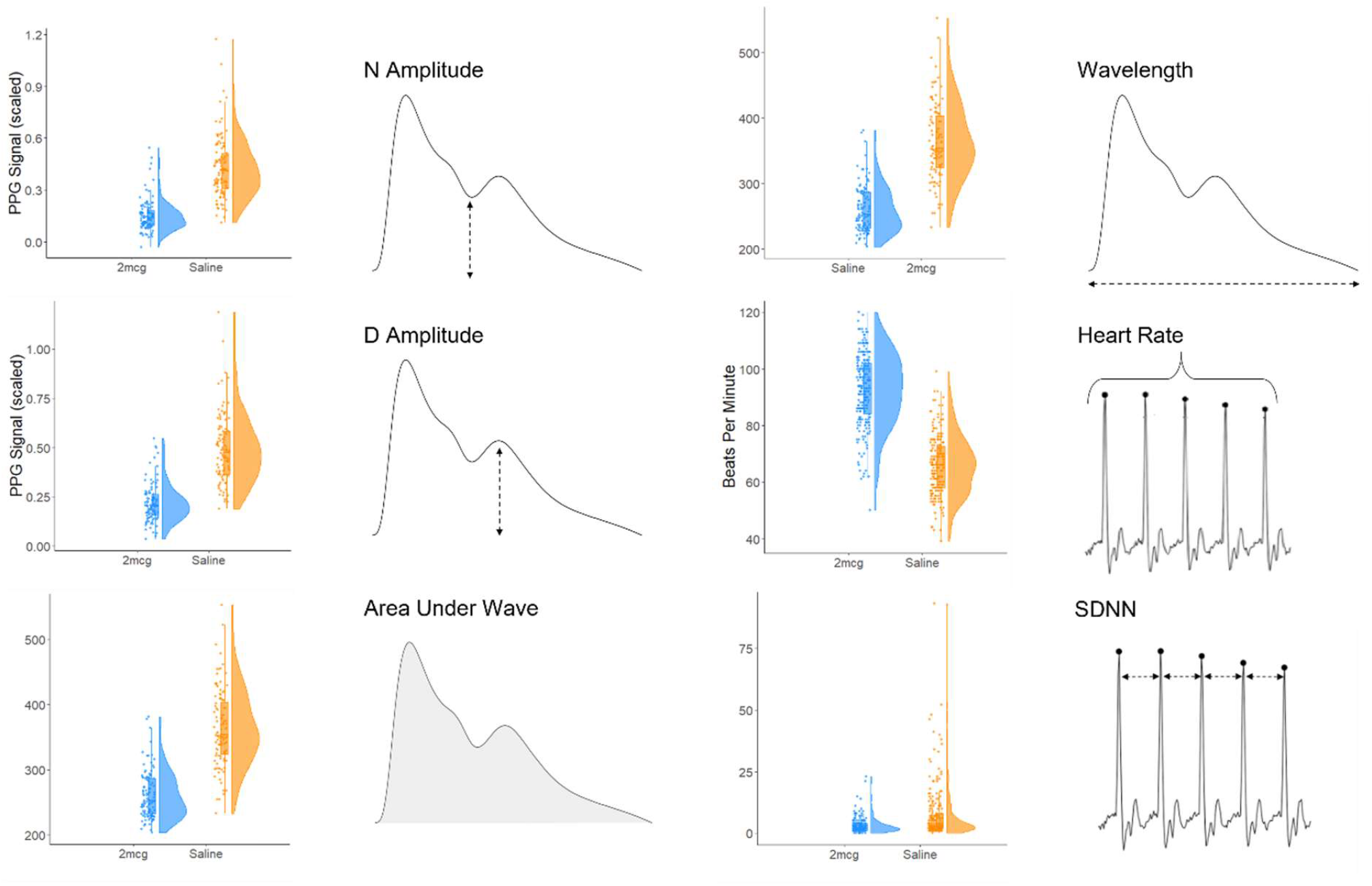
Standard features derived from the PPG signal (heart rate and SDNN as a measure of heart rate variability) and from pulse waveform (by area, or by identification of fiducial points; O, S, N, D) across 0 mcg and 2 mcg dose levels. Each point in the raincloud plots (Allen, et al. 2019) corresponds to the average (mean) value for a single participant at one dose level. A graphical representation of each feature indicating how they were measured is included (heart rate and SDNN were derived from inter-beat intervals of the PPG signal). See table 1 for accompanying statistics.

Figure 9 shows those model parameter outputs which characterized the difference in dose level most effectively, a combination of both excess and decay parameters (for remaining parameters see supplementary material (S2)). The systolic and second reflectance waves were modelled as wider at the 2 mcg level, whilst the first reflectance wave was highly variable in width and earlier in time. The 1st baseline was lower at 2 mcg, and the decay rate was increased. Both contributed to the modelling of a much steeper decay at 2 mcg than at saline.

**Table 1.** Statistical comparisons across isoproterenol and placebo conditions from standard and HED model-derived measures.

| **Standard Measures** | (see figure 8) |  |  | **HED Model** | (see figure 9) |  |  |
| --- | --- | --- | --- | --- | --- | --- | --- |
|  | Isoproterenol versus Saline |  |  |  | Isoproterenol versus Saline |  |  |
| **Parameter** | **t (df)** | **P value** | **Cohen's d** | **Parameter** | **t (df)** | **P value** | **Cohen's d** |
| N Amplitude | -13.8 (155) | <0.001 | 1.92 | S Width | 9.4 (203) | <0.001 | 1.3 |
| D Amplitude | -13.9 (162) | <0.001 | 1.93 | R1 Width | -8.6 (141) | <0.001 | 1.19 |
| Area Under Wave | -11.3 (144) | <0.001 | 1.57 | R2 Width | 7.1 (168) | <0.001 | 0.98 |
| Wavelength* | -15.5 (173) | <0.001 | 2.15 | R1 Time | -5.2 (202) | <0.001 | 0.73 |
| Heart Rate | 23.5 (401) | <0.001 | 2.29 | 1<sup>st</sup> Baseline | -8.04 (194) | <0.001 | 1.11 |
| Heart Rate Variability (SDNN) | -6 (239) | <0.001 | 0.59 | Decay Rate** | -17.3 (206) | <0.001 | 2.4 |

\* Best performing measure from standard analysis
\*\*Best performing parameter from HED model analysis

**Figure 9.**
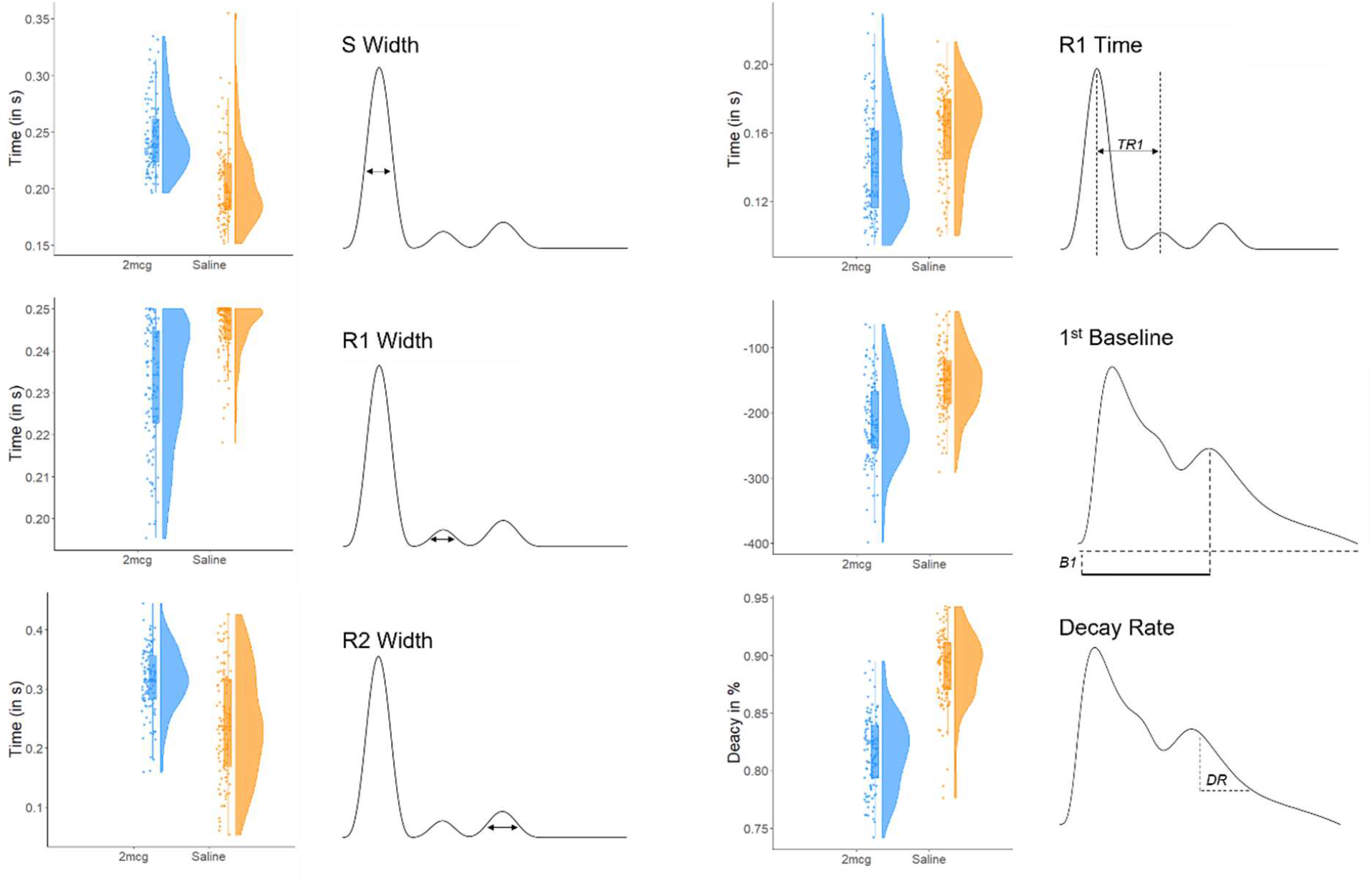
Novel features derived from the modelled waveform across the saline and 2 mcg dose levels. Each point in the raincloud plots (Allen, et al. 2019) corresponds to the average (mean) value for a single participant at one dose level. A graphical representation of each feature indicating how they were measured is included. See table 1 for accompanying statistics. (Note that the truncated distribution (especially notable for R1 width) is a result of a clamping algorithm applied to prevent over-fitting. (The algorithm was tending to use parameters for R1 (the first reflectance wave) to optimise overall fit). This was prevented by restricting permissible values).

In table 1, we summarise the statistics and effects sizes from each of the parameters shown in figures 8 and 9

ROC curves were generated to assess classifying performance of derived features, shown in figure 10 (a and b). The 5 features with the highest AUC values in 11a appeared to be similar in discriminative ability, which may indicate a redundancy between measures. Wavelength, a measure directly related to heart rate, performed best numerically, suggesting that for the purposes of classifying between stimulation with isoproterenol versus saline, more traditional morphological assessment waveform using traditional measures was of no added value. SDNN, the standard deviation of inter-beat intervals across the selected region of the time series, performed less well. This may be due to the variability in inter-beat interval due to the fact that we were confined to a limited number of beats relating to the duration of the infusion periods.

**Figure 10.**
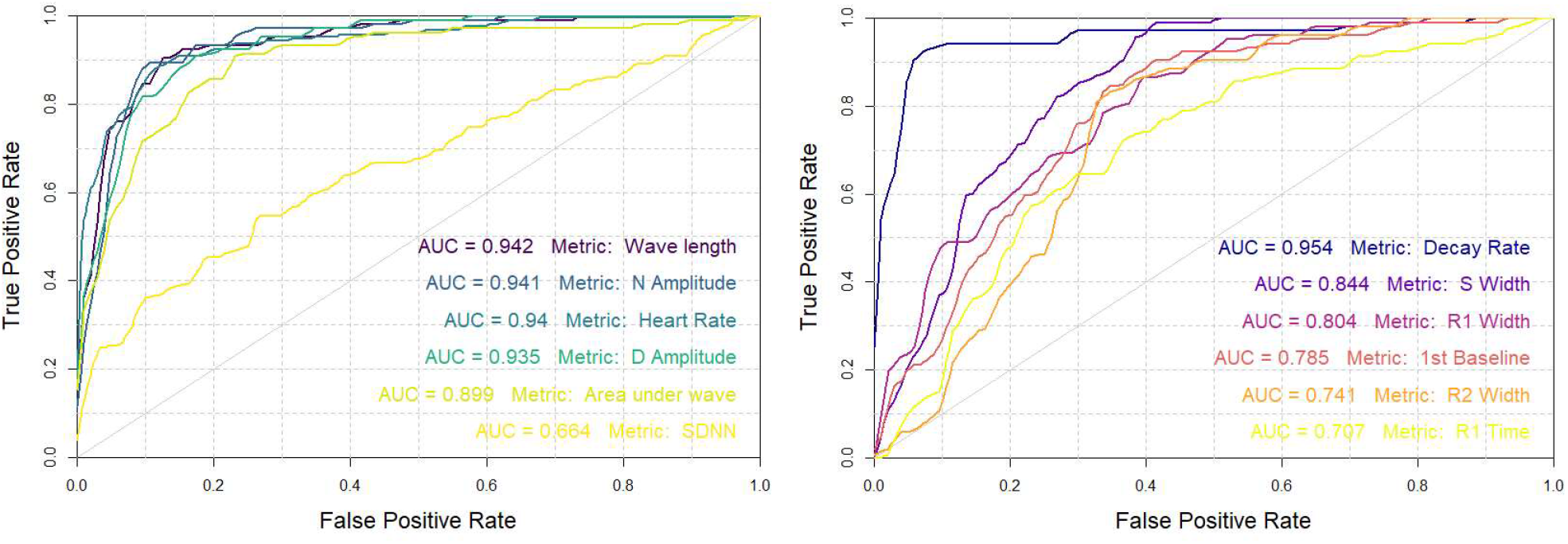
a: ROC curve of top performing (highest AUC) standard features, heart rate, and heart rate variability in classifying between saline and 2 mcg morphologies. b: ROC curve of top performing (highest AUC) HED model parameters (with best performing morphological feature, wavelength, overlaid in blue) in classifying between saline and 2 mcg morphologies.

Figure 10b shows the best performing model parameters. Decay rate showed the highest AUC value, with, notably, a higher numerical performance in distinguishing isoproterenol from saline than any of the more standard measures (shown in figure 10a) including wave length and heart rate. These findings suggest that the added granularity offered by the proposed model was useful and informative in this instance. This, however, is a qualitative observation and formal testing using Deong’s test for two correlated ROC curves was used to compare the “best-performing” measure from the HED model (decay rate) and that from more standard approaches (wavelength). These were not significantly different (p = 0.495; Z = 0.682).

## 4 DISCUSSION

We present a new model – the HED model – with an accompanying automated pipeline for analysing morphological characteristics of the PPG waveform. While standard approaches focus primarily on basic features such as HR and HRV, we argue that the waveform itself contains valuable information about underlying physiological states relevant to psychological and human neuroscientific research. Previous analyses of the waveform have used particular landmarks such as the onset, systolic peak, dicrotic notch and diastolic peak (*OSND*) to identify features of relevance to individual state and trait characteristics, including anthropometrics (Broyd *et al*., 2005), cardiovascular health (Girón, Millán and López, 2019) and current state of arousal (Arza *et al*., 2019). The HED model was motivated by this early promise and by the belief that a detailed, automated and physiologically-plausible approach to characterising the waveform more comprehensively will help to realise its potential. Below, we consider the HED model in relation to previous approaches to the PPG waveform, its biological plausibility and its sensitivity to a relevant pharmacological perturbation. We acknowledge that this model must be seen as early in development and consider its limitations before highlighting areas where we feel that the more comprehensive analysis that it will be of value to neuroscience and psychiatry.

By formally modelling, in addition to its component waves, a decay component, along with shifting baselines across the cardiac cycle, the model extends previous PDA approaches (Fleischhauer *et al*., 2020). It performed well on an external dataset and, moreover, this performance matched, and indeed, numerically exceeded in terms of its ROC curve, several standard metrics, including HR and HRV. This capacity to distinguish between signal acquired during a pharmacological challenge and that acquired in the same subjects during a saline control suggests that it may prove a useful complement to these existing measures. Indeed, it is notable that an isoprenaline dose of 2mcg produces a marked increase in HR (by an average of 26 beats per minute in the current dataset). When assessing more subtle experimental manipulations, such as the imposition of psychological stress, which produce much smaller – if any – differences in heart rate, the morphology of the pulse wave may become increasingly valuable. While its potential remains to be seen, it has already shown promise in its performance, relative to HRV, in distinguishing physiological changes induced by cognitive demand (Pelaez MDC 2019). Below, we consider more deeply, the implications of our findings and, more generally, the potential value of modelling the waveform in this way.

Previous models using PDA have converged on a three-component waveform. With the addition of exponential decay as a separable element, our model produces a characterisation that relates to a concept from the field of arterial haemodynamics. The hybrid reservoir-wave model (Wang *et al*., 2003) aims to account for aortic pressure waveform morphology by decomposing it into ‘reservoir’ and ‘excess’ pressures. Doing so enables both windkessel and wave reflectance properties of the arterial tree to be captured. According to the hybrid model, the reservoir pressure accounts for arterial wall expansion properties and is constant throughout the arterial tree, while excess pressure accounts for local changes in pressure due to propagating waves at the measurement site, including the initial systolic and ensuing reflectance waves. Though lacking widespread acceptance (Mynard and Smolich, 2017; Segers *et al*., 2017), there is growing evidence that the hybrid model’s parameters are of cardiovascular prognostic value (Hughes and Parker, 2020).

While our model shares features with the reservoir-wave model, it is important to note a key difference: it relates to a peripheral (i.e. finger) measurement. Despite evidence of some equivalence between peripheral arterial measures and aortic ones (Peng, 2018), the PPG signal measures blood volume, rather than pressure, and at low wavelengths of light may incorporate time-delayed signals from different tissue depths (Liu *et al*., 2019). Nonetheless, the conceptual similarity to a model that has proven useful in central arterial studies lends credence to the one presented here.

A key test lay in the application of the approach to an independent dataset and its capacity to distinguish isoproteronol from saline effects within this dataset. The average goodness of fit achieved by the model is comparable to that of other PDA models (Sorelli, Perrella and Bocchi, 2018) though a standardized means of comparison has not been established. However, fits were equivalent across both isoproterenol and saline allowing us to make direct comparisons across these states. Results of the ensuing analysis suggested that one model parameter, the decay rate, showed numerical superiority – in terms of ROC and effect size measures (see table 1) over traditional pulse-derived measures. Though it is important to note that it did not show statistical superiority, this finding nonetheless suggests that characterizing the peripheral pulse waveform more thoroughly can yield useful information over and above the typical PPG-derived measures of HR and HRV.

The two-element windkessel model (Westerhof, Lankhaar and Westerhof, 2009) provides a useful framework for contextualizing our finding of the numerical superiority obtained by including the decay rate parameter. It models the time duration of exponential pressure decay in diastole (RC time) as the product of arterial compliance (C) and peripheral resistance (R). The model can therefore characterize changes in either R or C in terms of RC time. Since exponential decay rate is inversely related to exponential decay time, it is plausible that our modelled decay rate approximates RC time (insofar as PPG decay mimics central pressure decay). With this framework, we can link the reduced peripheral resistance induced by isoproterenol (Smith HJ 1967) (Green 1977) to the change in waveform morphology effectively characterized by decay rate. Furthermore, if decay rate is a parameter sensitive to key features of cardiovascular state (R and C), then it may in turn be sensitive to measures related to these (e.g., cardiac output, pulse pressure, stroke volume). Thus, decay rate has plausible links to peripheral cardiovascular features beyond those directly pertaining to the heart, which may explain its excellent performance in distinguishing isoproteronol from saline infusion.

The model we propose, then, could be interpreted as following a simple underlying principle: An initial pressure wave, corresponding to cardiac systole, decays exponentially. The nature of this decay is determined both by arterial compliance and also by ensuing reflectance waves which are, themselves, influenced by compliance and a number of other features relating to an individual’s cardiovascular state. While we have examined the impact of a pharmacological perturbation within subjects, it will be interesting in the future to determine whether the model is sensitive to other variations across individuals and groups.

There are several limitations that should be considered. The data used in the development of the proposed model were obtained from two different hardware set-ups, both of which were transmission-based. Differing placement of PPG devices, and the use of reflectance rather than transmission-based measures will have an impact on the measured pulse waveforms. Whilst our model has demonstrated robustness to different morphologies of waveform, we cannot confirm its validity on sources such as these, from which different signals may arise. Related to this, the population of participants in the ISO dataset were predominantly younger and female, and there was also a range of psychiatric conditions represented. As with more widely-used measures such as HR, characteristic waveforms will likely differ across sex and age (which themselves may influence the effects of isoproterenol (Stratton 1999)) as well as other factors such as height, and it remains to be demonstrated that the model pipeline will generalise across these factors. Existing analyses of isolated waveform features have already identified features that differ with age (Allen and Murray 2002) or acute physiological stress (Celka P 2019) as well as cardiovascular (Elgendi et al 2019; Millasseau et al 2002; Lee et al 2011; Takazawa et al 1998) and mental (Kontaxis S 2021) health suggesting that this characterisation of the waveform across groups may prove informative.

In addition, it is likely that the performance of our model could be optimized further. When increasing degrees of freedom with additional parameters, the risk of overfitting should be considered. Standard deviations of waves fitted by the model indicated an increased within-participant variability in the R1 Time and R1 width parameters at the 2mcg dose level (see supplementary material – S5). On visual inspection of these time series, a marked variability in the beat-to-beat fitting of the 1st reflectance wave was noted in some cases, indicating a tendency to optimize fit at the expense of robustness. This behaviour only becomes apparent when the diastolic portion of the waveform has dropped to below a certain threshold, when the position of a 1st reflectance wave in the data becomes visually ambiguous. This limitation could be overcome by an initial calibration period at baseline when the parameters of R1 can be ascertained more reliably. Stricter constraints could then be applied to any periods of deviation from baseline. We suggest any future implementations of this model take such an approach to avoid similar instances of over-flexibility.

Nevertheless, we argue that there are several good reasons for pursuing a robust, automated approach to characterising the morphology of the pulse waveform generated by standard PPG hardware. The waveform is influenced by several underlying factors that are of interest in terms of physiology and, potentially, psychology. As such, we have the possibility of establishing a valuable complement to the more standard outputs of heart rate and heart rate variability.

Given that the waveform morphology reflects vascular characteristics which are the subject of several controlling factors, its study opens potentially important avenues of research. As an example, there has been increasing interest in the influence of the intra-cellular Adhesion Molecule-1 (ICAM) in a range of psychiatric disorders (Müller 2019) and given too that this molecule is an important regulator of vascular elasticity, it may be that the pulse waveform offers itself as a valuable marker for alterations in ICAM activity in certain psychiatric illnesses. The possibility that the waveform may signal subtle cardiovascular changes that occur in association with longer term changes in psychological health has already been examined in the context of, for example, depression (Dregan *et al*., 2020) and observations that other conditions, such as anorexia nervosa (AN) are associated with altered arterial stiffness (Tonhajzerova I 2020; Jenkins ZM 2021) hint at its potential usefulness in psychiatry. Indeed, features present in the waveform have been related to aspects of mental health in the case of depression, but cardiovascular changes are also well recognized hallmarks of other long term mental illnesses such as schizophrenia and AN. With the increasing prevalence of PPG-based wearables, the pulse waveform may offer a means of remotely monitoring physical and mental health (Myin-Germeys *et al*., 2018).

Related to the above, the growing field of interoceptive neuroscience and its focus on measures that characterize subjective and objective elements of brain-body interaction would appear to be complementary to the work presented here Critchley and Garfinkel, 2015; Khalsa et al., 2018; Allen et al. 2020; Allen 2020) Heartbeat detection tasks have been used frequently to measure interoceptive perceptual ability (Brener and Ring, 2016) since heartbeats represent a convenient interoceptive signal, and subjective perception of heartbeats reported by participants can be compared with objective measurement of these beats. However, PPG measures that go beyond simple indices of systole and diastole in relating interoceptive abilities to psychological states may be of value in developing our understanding of links between cardiovascular states and interoception (Schandry, Bestler and Montoya, 1993), including studies that employ peripheral perturbations to modulate pulsatility of the vascular tree in healthy (Khalsa, et al. 2009), psychiatric (Khalsa et al 2015) and neurological populations (Khalsa et al 2009), and those examining the accuracy and precision of metacognitive beliefs (Garfinkel et al 2016; Legrand et al 2021), possibly in combination. While the subjective experience that forms the basis for successful cardiac interoception (i.e. ‘cardioception’) is not clear, it is feasible that the pulsatility of the vascular tree contributes to it. As such, we should consider the possibility that individual variation in cardioceptive abilities does not rely solely on higher order sensitivity but also on the signal itself, with the nature of pulse wave morphology potentially contributing to this. Characterising wave differences across individuals will thus be crucial in relating cardiac interoceptive abilities to psychological states, traits, and psychiatric disorders. Moreover, relating characteristics of the waveform to the nature and magnitude of heartbeat evoked potentials (Pollatos and Schandry, 2010; Park and Blanke 2019; Coll et al 2021; Babo-Rebelo et al 2016) may prove useful in understanding generators and modulators of these potentials. Crucially, too, if the waveform differs in individuals with psychiatric illness, it may be that their altered interoceptive capacity is secondary to a deeper physiological alteration, which may explain the limited benefits of extant attempts at cardioceptive biofeedback training (Rominger C 2021). Future work should focus on validating the model with additional datasets and assessing its ability to characterize a range of psychophysiological states across individuals. Obtaining reliable signals of sufficient quality will be important for this. Reassuringly, this in itself is a field of great interest and innovation (McDuff *et al*., 2020). Moreover, existing early observations that features of the PPG waveform may outperform more standard measures like HRV (Pelaez MDC 2019) are promising in this regard. Thus, the more fully we understand the PPG signal, and how it varies across individuals, and across physiological and psychological states, the more realistically we might be able to understand interoceptive abilities and perhaps even develop a clearer idea of the causal chain between brain and body feedback interactions.

Overall, we argue that the shape of the pulse waveform, which has tended to be overlooked (Murray and Foster, 1996), contains information that will prove useful in a number of settings, including assessments of psychological states of arousal. We present the HED model, a robust automated model for characterising this waveform and show that its parameters compare favourably with more established ones such as heart rate and heart rate variability in distinguishing the effects of a sympathomimetic drug from a saline control. In this respect, we show that the waveform conveys valuable information on acute changes in cardiovascular state.

We suggest that the use of this theoretically-motivated model extends existing approaches using pulse decomposition. How the parameters of the waveform perform in distinguishing states that differ more subtly than the drug effects seen here remains to be seen. The fact that features of the wave are influenced by a number of relevant factors give some grounds for supposing that it may provide a rich and sensitive objective index of physiological or psychological arousal.

## Supporting information

Full supplemental material

## Acknowledgements

This work arose from a collaboration between Ninja Theory Ltd and University of Cambridge (details at theinsightproject.com). L D-W is supported by a generous donation from Ninja Theory Ltd to University of Cambridge. Funding was provided by the Bernard Wolfe Health Neuroscience Fund to PCF and a Wellcome Trust Investigator Award to PCF (Reference No. 206368/Z/17/Z). The work has been supported in part by the National Institute of Mental Health (grant K23MH112949 to Dr Khalsa); National Institute of General Medical Sciences (grant P20GM121312 to Drs. Khalsa and Paulus), and The William K. Warren Foundation. The content is solely the responsibility of the authors and does not necessarily represent the official views of the National Institutes of Health. MA is supported by a Lundbeckfonden Fellowship (under Grant [R272–2017-4345]), and the AIAS-COFUND II fellowship programme that is supported by the Marie Skłodowska-Curie actions under the European Union’s Horizon 2020 (under Grant [754513]), and the Aarhus University Research Foundation.

## References

Abbod, M. et al. (2011) ‘Developing a monitoring psychological stress index system via photoplethysmography’, Artificial Life and Robotics, 16(3), pp. 430–433. doi: 10.1007/s10015-011-0976-y.

Akl, T. J. et al. (2014) ‘Quantifying tissue mechanical properties using photoplethysmography’, Biomedical Optics Express, 5(7), pp. 2362–2375. doi: 10.1364/BOE.5.002362.

Allen, J, and A Murray. 2002. “Age-related changes in peripheral pulse timing characteristics at the ears, fingers and toes.” Journal of Human Hypertension 16: 711–717. doi:10.1038/sj.jhh.1001478.

Allen, J. (2007) ‘Photoplethysmography and its application in clinical physiological measurement’, Physiological Measurement, 28(3), pp. R1–R39. doi: 10.1088/0967-3334/28/3/R01.

Allen, M. 2020. “Unravelling the Neurobiology of Interoceptive Inference.” Trends in Cognitive Sciences 24 (4): 265–266. doi:10.1016/j.tics.2020.02.002.

Allen M, Legrand N, Costa Correa CM, and Fardo F. 2020. “Thinking through prior bodies: autonomic uncertainty and interoceptive self-inference.” Behavioral and Brain Sciences 43 (e91). doi:10.1017/S0140525X19002899.

Allen, M, Poggiali D, Whitaker K, Marshall TR, and Kievit RA. 2019. “Raincloud plots: a multi-platform tool for robust data visualization.” Wellcome Open Research 4 (63). doi:10.12688/wellcomeopenres.15191.1.

Arza, A. et al. (2019) ‘Measuring acute stress response through physiological signals: towards a quantitative assessment of stress’, Medical & Biological Engineering & Computing, 57(1), pp. 271–287. doi: 10.1007/s11517-018-1879-z.

Asekar, M. S. (2018) ‘Development of Portable Non-Invasive Blood Glucose Measuring Device Using NIR Spectroscopy’, in 2018 Second International Conference on Intelligent Computing and Control Systems (ICICCS). 2018 Second International Conference on Intelligent Computing and Control Systems (ICICCS), pp. 572–575. doi: 10.1109/ICCONS.2018.8663039.

Babo-Rebelo M, Richter CG, Tallon-Baudry C. 2016. “Neural Responses to Heartbeats in the Default Network Encode the Self in Spontaneous Thoughts.” Journal of Neuroscience 36 (30): 7829–7840. doi:10.1523/JNEUROSCI.0262-16.2016.

Baruch, M. C. et al. (2011) ‘Pulse Decomposition Analysis of the digital arterial pulse during hemorrhage simulation’, Nonlinear Biomedical Physics, 5, p. 1. doi: 10.1186/1753-4631-5-1.

Brener, J. and Ring, C. (2016) ‘Towards a psychophysics of interoceptive processes: the measurement of heartbeat detection’, Philosophical Transactions of the Royal Society B: Biological Sciences, 371(1708), p. 20160015. doi: 10.1098/rstb.2016.0015.

Broyd, C. et al. (2005) ‘Association of pulse waveform characteristics with birth weight in young adults’, Journal of Hypertension, 23(7), pp. 1391–1396. doi: 10.1097/01.hjh.0000173522.98728.58.

Cameron OG, Minoshima S. 2002. “Regional brain activation due to pharmacologically induced adrenergic interoceptive stimulation in humans.” Psychosomatic medicine 851–861. doi:10.1097/01.psy.0000038939.33335.32.

Celka P, Charlton PH, Farukh B, Chowienczyk P, Alastruey J. 2019. “Influence of mental stress on the pulse wave features of photoplethysmograms.” Healthcare Technology Letters 7 (1): 7–12. doi:10.1049/htl.2019.0001.

Charlton, P. H. et al. (2018) ‘Assessing mental stress from the photoplethysmogram: a numerical study’, Physiological Measurement, 39(5), p. 054001. doi: 10.1088/1361-6579/aabe6a.

Cleaveland CR, Rangno RE, Shand DG. 1972. “A standardized isoproterenol sensitivity test. The effects of sinus arrhythmia, atropine, and propranolol.” Archives of Internal Medicine 47–52. doi:10.1001/archinte.130.1.47.

Coll MP, Hobson H, Bird G, Murphy J. 2021. “Systematic review and meta-analysis of the relationship between the heartbeat-evoked potential and interoception.” Neuroscience and Biobehavioural Reviews 122: 190–200. doi:10.1016/j.neubiorev.2020.12.012.

Contrada RJ, Dimsdale J, Levy L, Weiss T. 1991. “Effects of isoproterenol on T-wave amplitude and heart rate: a dose-response study.” Psychophysiology 458–462. doi:10.1111/j.1469-8986.1991.tb00731.x.

Critchley, H. D. and Garfinkel, S. N. (2015) ‘Interactions between visceral afferent signaling and stimulus processing’, Frontiers in Neuroscience, 9. doi: 10.3389/fnins.2015.00286.

Dawber, T. R., Thomas, H. E. and McNamara, P. M. (1973) ‘Characteristics of the dicrotic notch of the arterial pulse wave in coronary heart disease’, Angiology, 24(4), pp. 244–255. doi: 10.1177/000331977302400407.

Domschke, K. et al. (2010) ‘Interoceptive sensitivity in anxiety and anxiety disorders: An overview and integration of neurobiological findings’, Clinical Psychology Review, 30(1), pp. 1–11. doi: 10.1016/j.cpr.2009.08.008.

Dregan, A. et al. (2020) ‘Associations Between Depression, Arterial Stiffness, and Metabolic Syndrome Among Adults in the UK Biobank Population Study: A Mediation Analysis’, JAMA psychiatry, 77(6), pp. 598–606. doi: 10.1001/jamapsychiatry.2019.4712.

Elgendi, M. (2012) ‘On the Analysis of Fingertip Photoplethysmogram Signals’, Current Cardiology Reviews, 8(1), pp. 14–25. doi: 10.2174/157340312801215782.

Elgendi, M. et al. (2019) ‘The use of photoplethysmography for assessing hypertension’, npj Digital Medicine, 2(1), pp. 1–11. doi: 10.1038/s41746-019-0136-7.

Elgendi, M., Liang, Y. and Ward, R. (2018) ‘Toward Generating More Diagnostic Features from Photoplethysmogram Waveforms’, Diseases (Basel, Switzerland), 6(1), p. E20. doi: 10.3390/diseases6010020.

Fleischhauer, V. et al. (2020) ‘Pulse decomposition analysis in photoplethysmography imaging’, Physiological Measurement, 41(9), p. 095009. doi: 10.1088/1361-6579/abb005.

Garfinkel SN, Manassei MF, Hamilton-Fletcher G, In den Bosch Y, Critchley HD, Engels M. 2016. “Interoceptive dimensions across cardiac and respiratory axes.” Philosophical Transactions of the Royal Society of London: B Biological Sciences 371 (1708). doi:10.1098/rstb.2016.0014

Girón, N. A., Millán, C. A. and López, D. M. (2019) ‘Systematic Review on Features Extracted from PPG Signals for the Detection of Atrial Fibrillation’, Studies in Health Technology and Informatics, 261, pp. 266–273.

glneurotech (2020) ‘BioRadio’, Great Lakes NeuroTechnologies, 11 March. Available at: https://www.glneurotech.com/products/bioradio/ (Accessed: 29 June 2021).

Green, J F. 1977. “Mechanism of action of isoproterenol on venous return.” American Journal of Physiology 232 (2): 152–156. doi:10.1152/ajpheart.1977.232.2.H152.

Hassanpour MS, Simmons WK, Feinstein JS, Luo Q, Lapidus RC, Bodurka J, Paulus MP, Khalsa SS. 2018. “The Insular Cortex Dynamically Maps Changes in Cardiorespiratory Interoception.” Neuropsychopharmacology 426–434. doi:10.1038/npp.2017.154.

Hughes, A. D. and Parker, K. H. (2020) ‘The modified arterial reservoir: An update with consideration of asymptotic pressure (P∞) and zero-flow pressure (Pzf)’, Proceedings of the Institution of Mechanical Engineers, Part H: Journal of Engineering in Medicine, 234(11), pp. 1288–1299. doi: 10.1177/0954411920917557.

Imholz, B. P. et al. (1998) ‘Fifteen years experience with finger arterial pressure monitoring: assessment of the technology’, Cardiovascular Research, 38(3), pp. 605–616. doi: 10.1016/s0008-6363(98)00067-4.

Jenkins ZM, Phillipou A, Castle DJ, Eikelis N, Lambert EA. 2021. “Arterial stiffness in underweight and weight-restored anorexia nervosa.” Psychophysiology. doi:10.1111/psyp.13913.

Khalsa SS, Craske MG, Li W, Vangala S, Strober M, Feusner JD. 2015. “Altered interoceptive awareness in anorexia nervosa: Effects of meal anticipation, consumption and bodily arousal.” International Journal of Eating Disorders 48 (7): 889–897. doi:10.1002/eat.22387.

Khalsa SS, Feinstein JS, Li W, Feusner JD, Adolphs R, Hurlemann R. 2016. “Panic Anxiety in Humans with Bilateral Amygdala Lesions: Pharmacological Induction via Cardiorespiratory Interoceptive Pathways.” Journal of Neuroscience 3559–3566. doi:10.1523/JNEUROSCI.4109-15.2016.

Khalsa SS, Rudrauf D, Damasio AR, Davidson RJ, Lutz A, Tranel D. 2008. “Interoceptive awareness in experienced meditators.” Psychophysiology 671–677. doi:10.1111/j.1469-8986.2008.00666.x.

Khalsa SS, Rudrauf D, Feinstein JS, Tranel D. 2009. “The pathways of interoceptive awareness.” Nature Neuroscience 12 (12): 1494–1496. doi:10.1038/nn.2411.

Khalsa SS, Rudrauf D, Sandesara C, Olshansky B, Tranel D. 2009. “Bolus isoproterenol infusions provide a reliable method for assessing interoceptive awareness.” International Journal of Psychophysiology 34–45. doi:10.1016/j.ijpsycho.2008.08.010

Khalsa, S. S. et al. (2018) ‘Interoception and Mental Health: A Roadmap’, Biological Psychiatry: Cognitive Neuroscience and Neuroimaging, 3(6), pp. 501–513. doi: 10.1016/j.bpsc.2017.12.004.

Khandoker, A. H. et al. (2017) ‘Suicidal Ideation Is Associated with Altered Variability of Fingertip Photo-Plethysmogram Signal in Depressed Patients’, Frontiers in Physiology, 8. doi: 10.3389/fphys.2017.00501.

Kontaxis S, Gil E, Marozas V, Lazaro J, Garcia E, Posadas-de Miguel M, Siddi S, Bernal ML, Aguilo J, Haro JM, de la Camara C, Laguna P, Bailon R. 2021. “Photoplethysmographic Waveform Analysis for Autonomic Reactivity Assessment in Depression.” IEEE Transactions on Biomedical Sciences 68 (4): 1271–2181. doi:10.1109/TBME.2020.3025908.

Legrand N, Nikolova N, Correa C, Braendholt M, Stuckert A, Kildahl N, Vejlo M, Fardo F, Allen M. 2021. “The heart rate discrimination task: a psychophysical method to estimate the accuracy and precision of interoceptive beliefs.” Preprint. https://www.biorxiv.org/content/10.1101/2021.02.18.431871v1.abstract.

Liu, J. et al. (2019) ‘Multi-Wavelength Photoplethysmography Enabling Continuous Blood Pressure Measurement With Compact Wearable Electronics’, IEEE Transactions on Biomedical Engineering, 66(6), pp. 1514–1525. doi: 10.1109/TBME.2018.2874957.

McDuff, D. et al. (2020) ‘Non-contact imaging of peripheral hemodynamics during cognitive and psychological stressors’, Scientific Reports, 10(1), p. 10884. doi: 10.1038/s41598-020-67647-6.

Millasseau, S. C. et al. (2002) ‘Determination of age-related increases in large artery stiffness by digital pulse contour analysis’, Clinical Science (London, England: 1979), 103(4), pp. 371–377. doi: 10.1042/cs1030371.

Millasseau, S. C. et al. (2006) ‘Contour analysis of the photoplethysmographic pulse measured at the finger’, p. 8.

Mulcahy, J. S. et al. (2019) ‘Heart rate variability as a biomarker in health and affective disorders: A perspective on neuroimaging studies’, NeuroImage, 202, p. 116072. doi: 10.1016/j.neuroimage.2019.116072.

Müller, N. (2019). The Role of Intercellular Adhesion Molecule-1 in the Pathogenesis of Psychiatric Disorders. Frontiers in Pharmacology. doi:10.3389/fphar.2019.01251

Lee, Q. Y. et al. (2011) ‘Multivariate classification of systemic vascular resistance using photoplethysmography’, Physiological Measurement, 32(8), pp. 1117–1132. doi: 10.1088/0967-3334/32/8/008.

Murray, W. B. and Foster, P. A. (1996) ‘The peripheral pulse wave: Information overlooked’, Journal of Clinical Monitoring, 12(5), pp. 365–377. doi: 10.1007/BF02077634.

Myin-Germeys, I. et al. (2018) ‘Experience sampling methodology in mental health research: new insights and technical developments’, World Psychiatry, 17(2), pp. 123–132. doi: 10.1002/wps.20513.

Mynard, J. P. and Smolich, J. J. (2017) ‘Wave potential: A unified model of arterial waves, reservoir phenomena and their interaction’, Artery Research, 18, pp. 55–63. doi: 10.1016/j.artres.2017.04.002.

Natarajan A, Pantelopoulos A, Emir-Farinas H, Natarajan P. 2020. “Heart rate variability with photoplethysmography in 8 million individuals: a cross-sectional study.” Lancet Digital health e650–e657. doi:10.1016/S2589-7500(20)30246-6

Nelder, J. A. and Mead, R. (1965) ‘A Simplex Method for Function Minimization’, The Computer Journal, 7(4), pp. 308–313. doi: 10.1093/comjnl/7.4.308.

Nitzan et al. (1994) ‘POWER SPECTRUM ANALYSIS OF SPONTANEOUS FLUCTUATIONS IN THE PHOTOPLETHYSMOGRAPHIC SIGNAL’, Journal of Basic and Clinical Physiology and Pharmacology, 5(3–4), pp. 269–276. doi: 10.1515/JBCPP.1994.5.3-4.269.

Park HD, Blanke O. 2019. “Heartbeat-evoked cortical responses: Underlying mechanisms, functional roles, and methodological considerations.” NeuroImage 197: 502–511. doi:10.1016/j.neuroimage.2019.04.081.

Parker, K. (2017) The reservoir-wave model⍰| Atlantis Press. Available at: https://www.atlantis-press.com/journals/artres/125924989 (Accessed: 28 June 2021).

Pelaez MDC, Albalate MTL, Sanz AH, Valles MA, Gil E. 2019. “Photoplethysmographic Waveform Versus Heart Rate Variability to Identify Low-Stress States: Attention Test.IEEE J Biomed Health Inform.” IEEE J Biomed Health Inform. 1940–1951. doi:10.1109/JBHI.2018.2882142.

Peng (2018) ‘Non-invasive measurement of reservoir pressure parameters from brachial-cuff blood pressure waveforms’. doi: 10.1111/jch.13411.

Penninx, B. W. J. H. (2017) ‘Depression and cardiovascular disease: Epidemiological evidence on their linking mechanisms’, Neuroscience and Biobehavioral Reviews, 74(Pt B), pp. 277–286. doi: 10.1016/j.neubiorev.2016.07.003.

Pollatos, O. and Schandry, R. (2010) ‘Accuracy of heartbeat perception is reflected in the amplitude of the heartbeat-evoked brain potential’, Psychophysiology, 41(3), pp. 476–482. doi: 10.1111/1469-8986.2004.00170.x.

Press, W. H. et al. (2007) Numerical Recipes 3rd Edition: The Art of Scientific Computing. 3rd edn. USA: Cambridge University Press.

Rinkevičius, M. et al. (2019) ‘Photoplethysmogram Signal Morphology-Based Stress Assessment’, in 2019 Computing in Cardiology (CinC). 2019 Computing in Cardiology (CinC), p. Page 1-Page 4. doi: 10.23919/CinC49843.2019.9005748.

Rominger C, Graßmann TM, Weber B, Schwerdtfeger AR. 2021. “Does contingent biofeedback improve cardiac interoception? A preregistered replication of Meyerholz, Irzinger, Withöft, Gerlach, and Pohl (2019) using the heartbeat discrimination task in a randomised control trial.” PLoS One 16 (3). doi:10.1371/journal.pone.0248246.

Sattar, R. R. M. et al. (2014) ‘Correlation between lipid profile and finger photoplethysmogram morphological properties among young men with cardiovascular risk: A preliminary result’, in 2014 IEEE Conference on Biomedical Engineering and Sciences (IECBES). 2014 IEEE Conference on Biomedical Engineering and Sciences (IECBES), pp. 602–606. doi: 10.1109/IECBES.2014.7047574.

Schandry, R., Bestler, M. and Montoya, P. (1993) ‘On the relation between cardiodynamics and heartbeat perception’, Psychophysiology, 30(5), pp. 467–474. doi: 10.1111/j.1469-8986.1993.tb02070.x.

Segers, P. et al. (2017) ‘Towards a consensus on the understanding and analysis of the pulse waveform: Results from the 2016 Workshop on Arterial Hemodynamics: Past, present and future’, Artery research, 18, pp. 75–80. doi: 10.1016/j.artres.2017.03.004.

Smith HJ, Oriol A, Morch J, McGregor M. 1967. “Hemodynamic studies in cardiogenic shock. Treatment with isoproterenol and metaraminol.” Circulation 35 (6): 1084–1091. doi:10.1161/01.cir.35.6.1084.

Smith R, Feinstein JS, Kuplicki R, Forthman KL, Stewart JL, Paulus MP, and Khalsa SS. Tulsa 1000 Investigators. 2021. “Perceptual insensitivity to the modulation of interoceptive signals in depression, anxiety, and substance use disorders.” Scientific reports. doi:10.1038/s41598-021-81307-3.

Sorelli, M., Perrella, A. and Bocchi, L. (2018) ‘Detecting Vascular Age Using the Analysis of Peripheral Pulse’, IEEE Transactions on Biomedical Engineering, 65(12), pp. 2742–2750. doi: 10.1109/TBME.2018.2814630.

Stratton, J R. 1999. “Effects of age and gender on the cardiovascular responses to isoproterenol.” The Journals of Gerontology: Series A 54 (9): 393–400. doi:10.1093/gerona/54.9.b401

Sviridova, N. and Sakai, K. (2015) ‘Human photoplethysmogram: new insight into chaotic characteristics’, Chaos, Solitons & Fractals, 77, pp. 53–63. doi: 10.1016/j.chaos.2015.05.005.

Takazawa, K. et al. (1998) ‘Assessment of Vasoactive Agents and Vascular Aging by the Second Derivative of Photoplethysmogram Waveform’, Hypertension, 32(2), pp. 365–370. doi: 10.1161/01.HYP.32.2.365.

Tigges, T. et al. (2017) ‘Model selection for the Pulse Decomposition Analysis of fingertip photoplethysmograms’, in 2017 39th Annual International Conference of the IEEE Engineering in Medicine and Biology Society (EMBC). 2017 39th Annual International Conference of the IEEE Engineering in Medicine and Biology Society (EMBC), pp. 4014–4017. doi: 10.1109/EMBC.2017.8037736.

Tonhajzerova I, Mestanikova A, Jurko A Jr, Grendar M, Langer P, Ondrejka I, Jurko T, Hrtanek I, Cesnekova D, Mestanik M. 2020. “Arterial stiffness and haemodynamic regulation in adolescent anorexia nervosa versus obesity.” Applied Physiology, Nutrition and Metabolism 81–90. doi:10.1139/apnm-2018-0867.

Wang, J.-J. et al. (2003) ‘Time-domain representation of ventricular-arterial coupling as a windkessel and wave system’, American Journal of Physiology-Heart and Circulatory Physiology, 284(4), pp. H1358–H1368. doi: 10.1152/ajpheart.00175.2002.

Wang, L. et al. (2013) ‘Multi-Gaussian fitting for pulse waveform using Weighted Least Squares and multi-criteria decision making method’, Computers in Biology and Medicine, 43(11), pp. 1661–1672. doi: 10.1016/j.compbiomed.2013.08.004.

Westerhof, B. E. et al. (2008) ‘Location of a reflection site is elusive: consequences for the calculation of aortic pulse wave velocity’, Hypertension (Dallas, Tex.: 1979), 52(3), pp. 478–483. doi: 10.1161/HYPERTENSIONAHA.108.116525.

Westerhof, N., Lankhaar, J.-W. and Westerhof, B. E. (2009) ‘The arterial Windkessel’, Medical & Biological Engineering & Computing, 47(2), pp. 131–141. doi: 10.1007/s11517-008-0359-2.

Zhang, O. et al. (2021) ‘Explainability Metrics of Deep Convolutional Networks for Photoplethysmography Quality Assessment’, IEEE access□: practical innovations, open solutions, 9, pp. 29736–29745. doi: 10.1109/access.2021.3054613.

Zhang Y, Weaver RG, Armstrong B, Burkart S, Zhang S, Beets MW. 2020. “Validity of Wrist-Worn photoplethysmography devices to measure heart rate: A systematic review and meta-analysis.” Journal of Sports Sciences 2021–2034. doi:10.1080/02640414.2020.1767348

