## Supplementary material for "Characterising the Photoplethysmography Pulse Waveform for Use in Human Neuroscience: The Hybrid Excess and Decay (HED) Model": Full supplemental material

Contents:

- S1: Automated Peak Detection
- S2: Morphological features and their calculation
- S3: Goodness of fit calculations and model performance (quantified by *a*NRMSE)
- S4: Identifying subset of heartbeats for modelling effects of isoproterenol and saline
- S5: Variability in Model Fitting

S1 – Automated Peak Detection

Peaks in PPG time series are more difficult to detect than their ECG counterparts, in part because of the large variations in signal between and within individuals. Peaks were most apparent in the first derivative of the signal but were not reliably detected with the relevant function (“findpeaks”) in a standard package (R).

A customised peak detection algorithm was therefore developed. The algorithm identifies peaks (denoted *w*) in the first derivative of the time series*.* An adaptive rolling window is employed to identify successive peaks in the time series. Within the window, inflection points are confirmed as peaks if they exceed thresholds relative to values within the window and across the entire time series. The size of the window is determined from previous inter-beat intervals and is therefore adaptive to changes in heart rate. It is also adaptive in its ability to skip forward (over artefactual regions) or widen in the forward direction (if inadequate beats detected). Artefacts are also detected by the window, again according to local and global thresholds.

Key assumptions made by the algorithm are as follows:

i (i) The duration of an inter-beat interval can be reasonably approximated from the previous inter-beat interval.
ii (ii) All peaks are inflection points in the data above the 95th percentile for amplitude within the window.
iii (iii) Peaks with extreme amplitude values relative to adjacent peaks and the entire time series should be labelled as artefactual.
iv (iv) Extended windows with no identifiable peaks are likely low-amplitude artefactual regions.
v (v) Peaks immediately preceding / following artefacts are likely to also be compromised and should therefore be removed. Our algorithm is currently removing 3 peaks either side of a detected artefact.
vi The first and last beat are excluded by default as they are often partial beats.

The algorithm’s performance was tested on a subset of 35 participants (70 time series) from an external dataset (see section 3.3).. A sensitivity of 0.97 and positive predictive value of 0.99 were achieved.

S3 - Morphological features and their calculation:

Morphological Features derived from each waveform are outlined below. See figure 1 in main text for referents.

| **Feature** | **Calculation** |
| --- | --- |
| Peak to Peak Time^4^ | $D_{x} - S_{x}$ |
| Notch to Peak Ratio | $\frac{N_{y}}{S_{y}}$ |
| Augmentation Index^5^ (also Reflection Index^6^) | $\frac{D_{y}}{S_{y}}$ |
| Crest Time^7^ | $S_{x} - O_{x}$ |
| Wavelength | $\lambda$  Time duration of entire waveform |
| Peak to Notch Time | $\frac{N_{x}- S_{x}}{\lambda}$ |
| Max Amplitude | Maximal amplitude of the waveform |
| Area Under Waveform^8^ | Integral of waveform |
| Inflection point area Ratio^9^ | $\frac{Area prior to N}{Area after N}$ |

*Morphological features derived from each waveform. Subscript notations of ‘x’ and ‘y’ refer to the x or y coordinates of the corresponding fiducial point, respectively. When scaling for amplitude, notch to peak ratio and augmentation index can be considered equivalent to N amplitude and D amplitude respectively.*

S3 – Goodness of fit calculation and model performance:

Calculation of goodness of fit as reported by Sorelli et al^2^:

| $NRMSE= 1-\left[ \sum_{i} \left( y_{i}- \hat{y_{i}} \right)^{2}- \sum_{i} \left( y_{i}- \bar{y} \right)^{2} \right]$ | $y$ = data value |
| --- | --- |
|  | $\hat{y}$ = modelled values |
|  | $\bar{y}$ = mean of data values |

Alternative goodness of fit measure, *aNRMSE,* as reported by Wang et al^3^:

| $aNRMSE = \frac{Sum of Squared residuals}{Sum of Squared data values}$ | $= \frac{\sum_{i} \left( y_{i}- \hat{y_{i}} \right)^{2}}{\sum_{i} \left( {y_{i}}^{2} \right)}*100$ | $y$ = data value |
| --- | --- | --- |
|  |  | $\hat{y}$ = modelled values |

The calculated *aNRMSE* value is an expression of the residual error as a percentage of the data. Values less than 2% are generally considered acceptable. Model performance on this measure is shown below (figure S1). The median aNRMSE value was 0.33%/

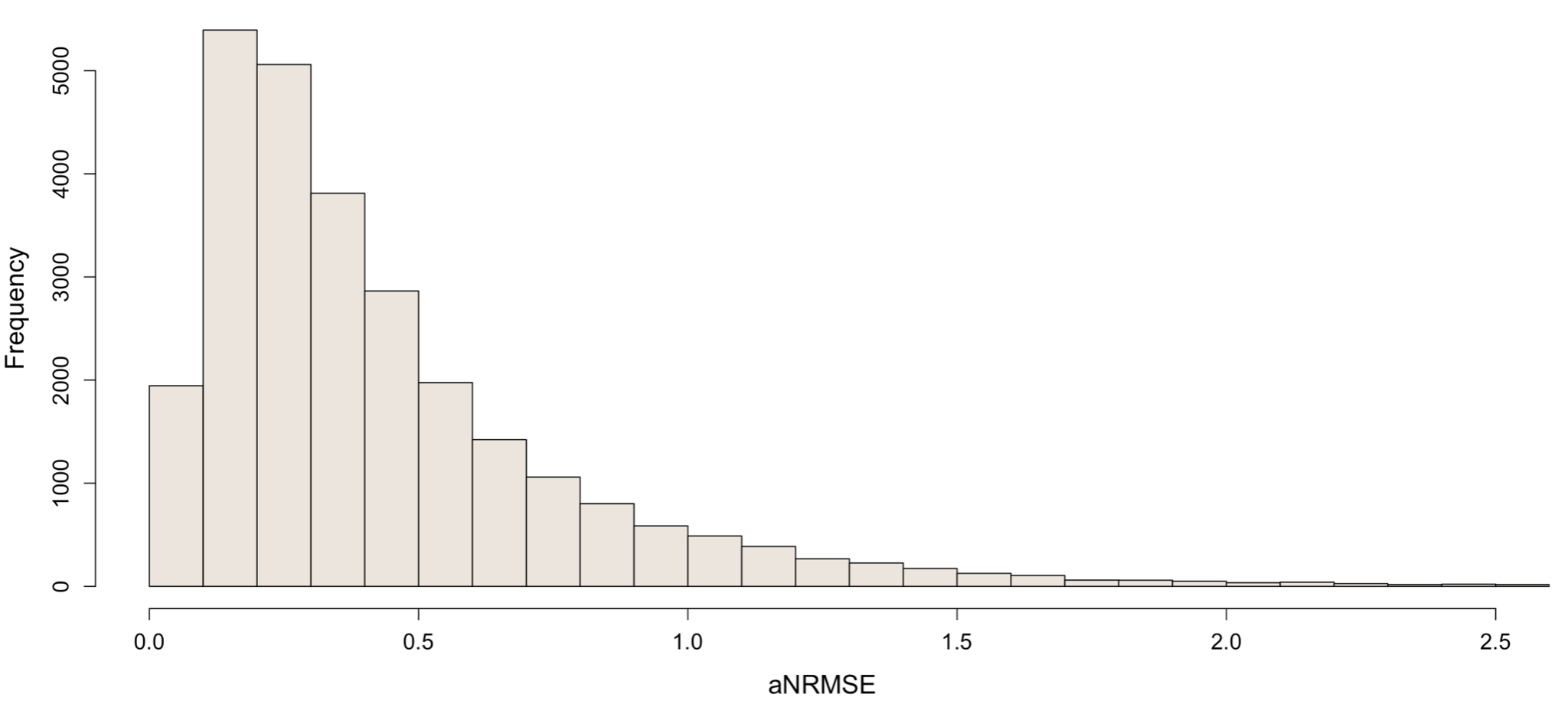

***Figure S1:*** *Distribution of aNRMSE values for all waves fitted by the model (bin width = 0.1).*

S4 – Identifying subset of heartbeats for modelling effects of isoproterenol and saline

The duration of effect of bolus isoprenaline infusion does not exceed 120 seconds^1^. Accordingly, time series were subsetted such that only beats within the timeframe corresponding to isoprenalines effect were analyzed. For each isoprenaline time series, inter-beat intervals (IBIs) were calculated, and a minimum inter-beat interval was identified. This minimum was taken as the peak of isoprenaline’s effect. The average inter-beat interval of the pre-infusion period (50 beats) was also calculated and taken as a baseline. From these values, a half-minimum was calculated. The closest IBI values to the half-minimum, before and after the minimum, were then selected as boundaries within which beats were subsetted (see figure S1). In order to maintain similar sample sizes across time series of different dose levels, the number of beats subsetted and lower boundary of the subsetting window were carried over from 2 mcg to 0 mcg time series.

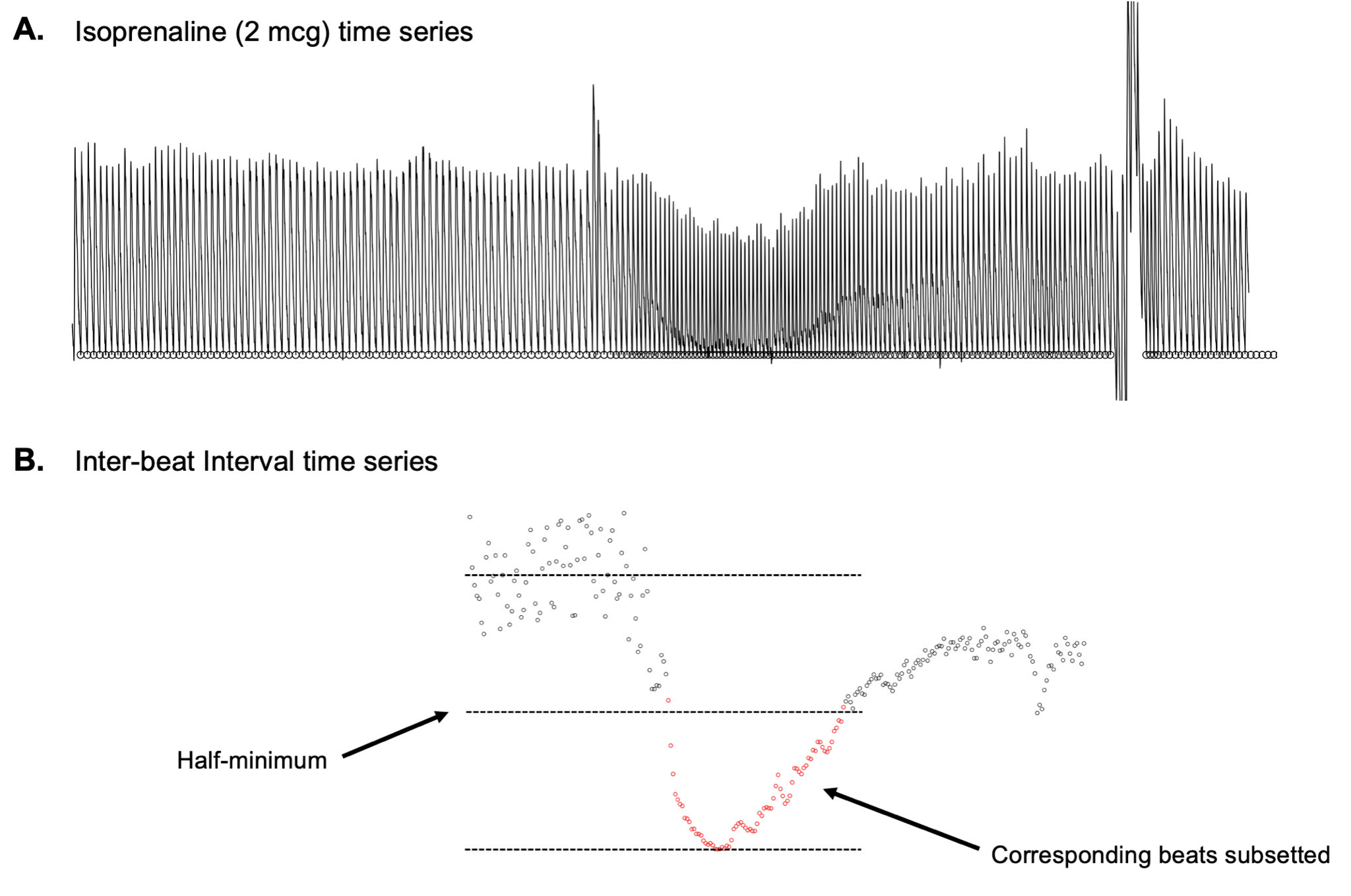

***Figure S2:*** ***A****: a typical time series during the infusion of isoprenaline (2 mcg). The change in morphology caused by isoprenaline is apparent.* ***B****: the inter-beat interval time series corresponding to* ***A****. Inter-beat intervals corresponding to the half-minimum if isoprenaline’s effect on heart rate are used to subset the time series.*

S6: Variability in Fitting:

Figure S3 shows a series of consecutive waves during the saline infusion. Despite a consistent appearance to the recorded waveforms, there is marked variability in beat-to-beat fitting for several parameters. This indicates a tendency for the model to optimize fit at the expense of robustness.

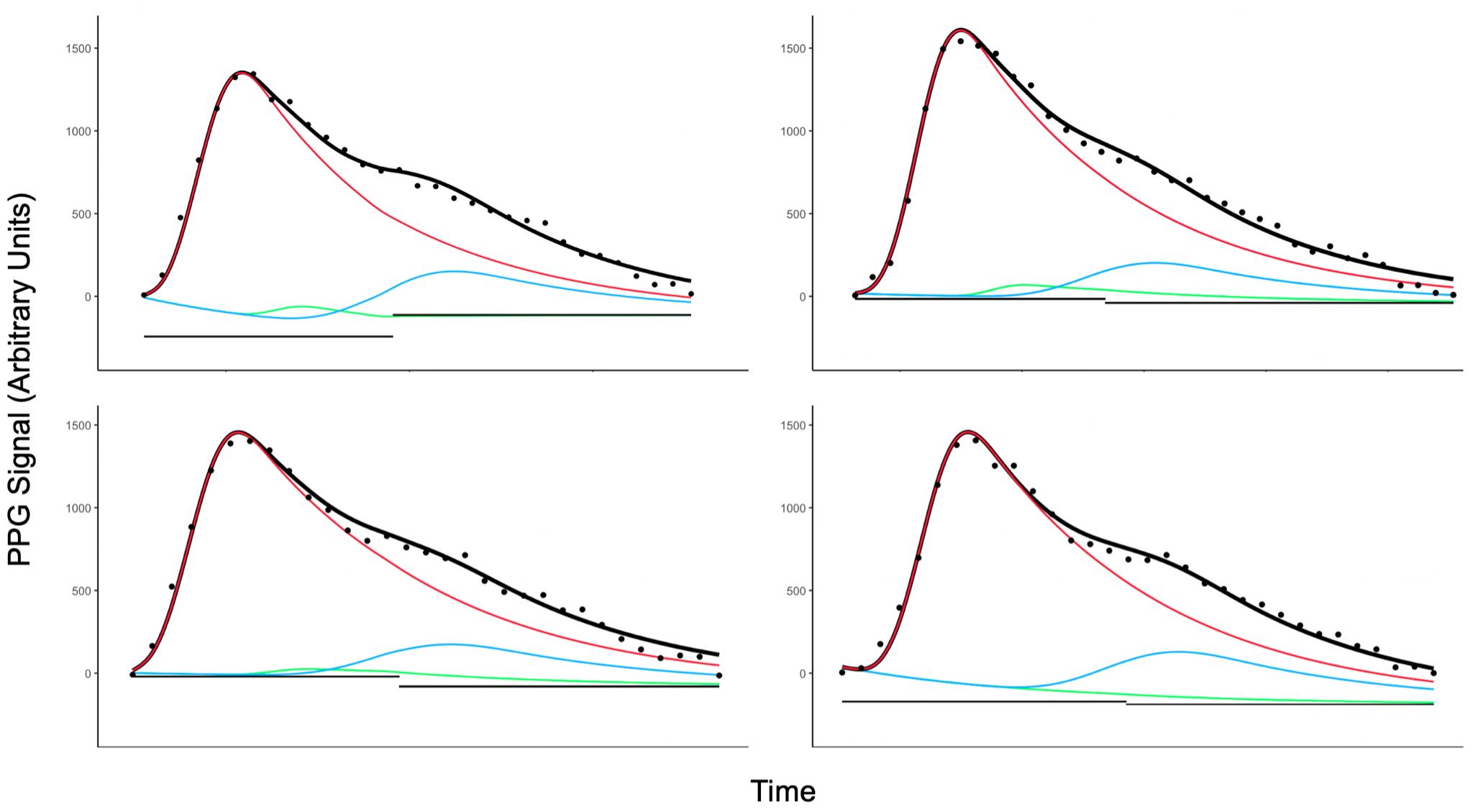

***Figure S3:*** *Variability in the fitting of waves within the ISO dataset. Despite similar morphologies, the variability in how consecutive waves are fitted is apparent – most notable are the baselines and first reflectance wave (note timing, width and decay rate parameters are fixed).*

**References:**

1. Khalsa, S. S. et al. Panic Anxiety in Humans with Bilateral Amygdala Lesions: Pharmacological Induction via Cardiorespiratory Interoceptive Pathways. J. Neurosci. **36**, 3559–3566 (2016).
2. Sorelli, M., Perrella, A. & Bocchi, L. Detecting Vascular Age Using the Analysis of Peripheral Pulse. IEEE Trans. Biomed. Eng. **65**, 2742–2750 (2018).
3. Wang, L., Xu, L., Feng, S., Meng, M. Q.-H. & Wang, K. Multi-Gaussian fitting for pulse waveform using Weighted Least Squares and multi-criteria decision making method. Comput. Biol. Med. **43**, 1661–1672 (2013).
4. Millasseau, S. C., Kelly, R. P., Ritter, J. M. & Chowienczyk, P. J. Determination of age- related increases in large artery stiffness by digital pulse contour analysis. Clin. Sci. Lond. Engl. 1979 **103**, 371–377 (2002).
5. Elgendi, M. On the Analysis of Fingertip Photoplethysmogram Signals. Curr. Cardiol. Rev. **8**, 14–25 (2012).
6. Padilla, J. M. et al. Assessment of relationships between blood pressure, pulse wave velocity and digital volume pulse. in 2006 Computers in Cardiology 893–896 (2006).
7. Alty, S. R., Angarita-Jaimes, N., Millasseau, S. C. & Chowienczyk, P. J. Predicting Arterial Stiffness From the Digital Volume Pulse Waveform. IEEE Trans. Biomed. Eng. **54**, 2268–2275 (2007).
8. Seitsonen, E. R. J. et al. EEG spectral entropy, heart rate, photoplethysmography and motor responses to skin incision during sevoflurane anaesthesia. Acta Anaesthesiol. Scand. **49**, 284–292 (2005).
9. Wang, L., Pickwell-MacPherson, E., Liang, Y. P. & Zhang, Y. T. Noninvasive cardiac output estimation using a novel photoplethysmogram index. in 2009 Annual International Conference of the IEEE Engineering in Medicine and Biology Society 1746–1749 (2009). doi:10.1109/IEMBS.2009.5333091.
